# New histone deposition recruits the DNA methylation maintenance machinery at sites of DNA damage repair

**DOI:** 10.64898/2026.07.27.740930

**Authors:** Margherita Mori, Sandra Piquet, Léa Girard, Laure Ferry, Kosuke Yamaguchi, Melina Farshchi, Elouan Bethuel, Olivier Kirsh, Magali Hennion, Pierre-Antoine Defossez, Sophie Polo

**Affiliations:** Laboratory of Epigenome Integrity, Epigenetics & Cell Fate Centre, UMR7216 CNRS, Université Paris Cité, Paris, 75013, France; Laboratory of Dynamics and Interpretation of DNA methylation, Epigenetics & Cell Fate Centre, UMR7216 CNRS, Université Paris Cité, Paris, 75013, France; Department of Chromosome Science, National Institute of Genetics, Research Organization of Information and Systems (ROIS), Mishima, 411-8540, Japan; BiBs Platform, Epigenetics & Cell Fate Centre, UMR7216 CNRS, Université Paris Cité, Paris, 75013, France

## Abstract

Faithful inheritance of DNA methylation contributes to the memory of epigenetic states and protects against disease. While the mechanisms underlying DNA methylation maintenance at replication forks are well characterized, whether and how DNA methylation is altered or maintained at sites of DNA damage repair is still poorly understood. Here, by exploiting sequencing, imaging and proteomic approaches in mammalian cells exposed to UV radiation, we show that the majority of DNA methylation marks are maintained during UV damage repair and we dissect the molecular machinery involved in DNA methylation control. We detect the recruitment to sites of repair synthesis of the DNMT1 and DNMT3A DNA methylating enzymes, driven by the DNMT1 cofactor UHRF1 and by UV damage repair endonucleases. We also uncover a crosstalk with histone dynamics, whereby newly deposited H3.3 histones at UV damage sites promote the recruitment of DNMT1. Functionally, we reveal the importance of the DNA methylation maintenance machinery for the transcriptional response to UV damage and sustained cell proliferation. This work provides a comprehensive picture of DNA methylation control mechanisms following DNA damage, with important implications for our understanding of human diseases with an altered methylome.

## INTRODUCTION

DNA methylation is an abundant DNA modification essential for cell and organism viability, which consists in the covalent linkage of a methyl group to the fifth carbon of the cytosine ring (5-methylcytosine, 5mC), and in the mammalian genome it occurs almost exclusively at cytosine followed by a guanine (CpGs) [1,2]. Faithfully inherited through cell divisions, DNA methylation is a key epigenetic mark that controls gene expression, represses repetitive elements, and, along with other chromatin marks, orchestrates cell identity [3,4]. DNA methylation is therefore tightly and dynamically controlled, both during embryonic development and throughout the cell cycle, and is established, maintained and erased by specific factors [5].

Deposited on previously unmethylated DNA by de novo DNA methyltransferases 3A and 3B (DNMT3A and DNMT3B) [6,7], DNA methylation is perpetuated during DNA replication by the maintenance DNA methyltransferase 1 (DNMT1), which copies parental methylation marks on the newly synthesized strand [8] with the help of the co-factor Ubiquitin-like with PHD and RING finger domains 1 (UHRF1). UHRF1 recruits and activates DNMT1 on hemi-methylated DNA through the deposition of ubiquitin groups on histone H3 and on the replication factor PCNA-Associated Factor 15 (PAF15), which are then recognized by DNMT1 [9,10]. UHRF1 itself binds methylated Ligase 1 (LIG1), and this protein modification is required for DNA methylation maintenance at the replication fork [11].

In the absence of functional DNA methylation maintenance, DNA methylation levels are diluted during successive rounds of replication, leading to passive demethylation [1]. DNA methylation is also actively erased by Ten-eleven translocation (TET) oxidative enzymes, which, through a series of oxidized intermediates, transform 5mC into a non-methylated cytosine [12]. The first and most abundant intermediate of 5mC demethylation is 5-hydroxy-methylcytosine (5hmC), which shows a widespread distribution in the mammalian genome [13]. While originally considered as a mere demethylation intermediate, 5hmC is a stable and functional DNA modification that plays a major role in transcription regulation [14,15].

Proper control of DNA methylation levels is crucial for preserving genome stability, physiological gene expression and cell identity, and DNA methylation alterations are widespread in human diseases including cancer and in aging [16–18]. Faithful inheritance of DNA methylation marks thus contributes to the memory of epigenetic states and protects against pathological development.

However, epigenome stability is challenged during all DNA transactions, in particular during the repair of damaged DNA [19]. Similar to DNA replication, DNA repair involves a DNA neo-synthesis step, with a need to re-establish DNA methylation patterns. While the molecular mechanisms underlying DNA methylation maintenance at replication forks are well characterized [5,20], a clear picture of DNA methylation control and dynamics at sites of DNA damage repair is still missing.

DNMT1 recruitment has been detected at DNA double-strand breaks (DSBs) and at sites of oxidative damage in mammalian cells [21–27]. In some instances, DNMT1 is accompanied by DNMT3A or DNMT3B [24,26], suggesting that these DNMTs may jointly contribute to methylating newly synthesized DNA during DNA damage repair. Supporting this possibility, DNMT3A was shown to cooperate with DNMT1 and UHRF1 to methylate a reporter gene during DSB repair in human cells, leading to a persistent heritable increase of DNA methylation at the repaired locus in a subset of cells [24]. Whether these factors similarly promote DNA methylation during the repair of endogenous loci is still unknown. UHRF1 accumulates at DSBs and interstrand crosslinks at endogenous loci and promotes their repair but whether and how this function coordinates with a role in DNA methylation maintenance is still elusive [28–32].

Whether DNA methylation marks are perfectly re-established upon DNA damage and repair is also debated. Both 5mC and 5hmC alterations have been detected in response to different types of DNA damage and with various spatial distributions and time frames. Several studies indeed report an increase of 5hmC mediated by TET enzymes upon DNA break induction, replication stress or UV irradiation in mammalian cells, either pan-nuclear [33,34] or localized at damage sites [35]. An increase in 5mC has been observed in response to DSBs and oxidative damage and contrasting 5mC alterations have been reported post UV damage [23,27,33,36,37].

Most studies on DNA methylation following DNA damage in mammalian cells have been performed upon induction of DSBs, which are repaired by different pathways according to the cell cycle stage and the chromatin context [38], and therefore are not a straightforward model to grasp DNA methylation dynamics at damage sites.

For a comprehensive dissection of DNA methylation dynamics in a more tractable DNA damage model, we decided to employ ultraviolet (UV) radiation, which is one of the most common environmental hazards and generates pyrimidine dimers on DNA [39]. Notably, cyclobutane pyrimidine dimers (CPDs), the most abundant UV photoproducts, form preferentially at methylated cytosines with an influence of the flanking sequence [40]. In this study we exploit UVC radiation, which activates exclusively the Nucleotide Excision Repair pathway (NER). The NER pathway repairs the broadest range of DNA lesions and does so throughout the cell cycle and in various chromatin contexts [41,42]. Remarkably, several NER factors modulate DNA methylation states even in undamaged cells. In plants, the damage sensor DNA damage binding protein 2 (DDB2) shapes DNA methylation patterns by controlling DNA methylation and demethylation pathways, and also prevents excessive changes in the DNA methylome after UV damage [43,44]. In mammalian cells, Xeroderma Pigmentosum, complementation group C (XPC), another factor involved in UV damage detection, promotes active DNA demethylation, of particular relevance during cellular reprogramming [45]. These findings suggest a possible involvement of the NER pathway in regulating DNA (de)methylation during DNA damage repair.

In this study, we characterize DNA methylation dynamics in response to UV damage in mammalian cells, and we decipher a machinery recruited to damage sites to control 5mC levels. We also uncover a dependency of this machinery both on NER and on new histone deposition at sites of DNA damage and reveal its functional relevance for the cellular response to UV damage.

## MATERIALS AND METHODS

### Cell culture and drug treatment

U2OS (ATCC HTB-96, human osteosarcoma, female), and NIH/3T3 GFP-DDB2 cells (ATCC CRL-1658, mouse embryonic fibroblasts, male, stably expressing a GFP-DDB2 transgene) [46] were grown in culture dishes (Falcon) at 37°C and 5% CO_2_ in Dulbecco’s Modified Eagle’s Medium (DMEM, Life Technologies) supplemented with 10% foetal bovine serum (EUROBIO) and antibiotics (100 U/ml penicillin and 100 μg/ml streptomycin, Life Technologies) and 200 µg/ml Hygromycin (Euromedex) for the NIH/3T3 GFP-DDB2 cell line. Before plating NIH/3T3 GFP-DDB2 cells on coverslips, the coverslips were treated with 20 μg/ml Collagen Type I (MERCK Millipore) and 2 μg/ml fibronectin (Sigma-Aldrich) in Phosphate Buffer Saline (PBS) for 1 h at 37°C to increase cell adhesion.

HCT116-LIG1-mClover-AID2 [47], HCT116-DNMT1-mClover-AID2 and HCT116-UHRF1-mClover-AID2 cell lines [48] were cultured in McCoy’s 5A medium (Sigma-Aldrich), supplemented with 10% FBS (EUROBIO), 2 mM L-glutamine and antibiotics (100 U/ml penicillin and 100 μg/ml streptomycin, Life Technologies) [49]. LIG1, DNMT1 and UHRF1 degradation was achieved after 2 h treatment with 1 μM auxin analogue (5-Ph-IAA, BioAcademia).

5-aza-2’-deoxycytidine (decitabine, Sigma-Aldrich) was used at 50 μM final concentration and the DNMT1 inhibitor GSK-3484862 (MedChem Express) was used at 4 μM final concentration for the indicated times. The G9a/GLP inhibitor UNC0642 (Sigma-Aldrich) was used at 1 μM for 72 h.

### Cell transfection

siRNAs (small interfering RNAs, **Supplementary Table S1**) purchased from Eurofins or Sigma-Aldrich were transfected into cells using Lipofectamine RNAiMAX (Invitrogen) following manufacturer’s instructions. Cell suspensions were counted with the automated cell counter Fluidlab R-300 (Anvajo). Cells were plated at 4x10^4^ cells/ml one day before transfection. The final concentration of siRNA in the culture medium was 50 nM. Cells were harvested 72 h post transfection.

For plasmid DNA transfection, cells were plated at 1x10^5^ cells/ml for NIH/3T3 GFP-DDB2 and 6x10^4^ cells/mL for U2OS one day before transfection and transfected with plasmid DNA (1 μg/ml final, **Supplementary Table S2**) using the X-treme GENE transfection reagent (Roche) according to manufacturer’s instructions.

### UVC lamp irradiation

Cells were irradiated with UVC (254 nm) using a low-pressure mercury lamp (Vilber Lourmat). Effective irradiation doses were determined using a VLX-3W dosimeter (Vilber Lourmat). For global UVC irradiation, cells on plates in Phosphate Buffer Saline (PBS) were exposed to 10 J/m^2^ UVC unless stated otherwise.

For local irradiation, cells grown on glass coverslips (12 mm diameter, type 1.5 thickness, Thorlabs) were irradiated through a polycarbonate filter (5 μm pore size, Millipore) with 500 J/m^2^ UVC. This local irradiation generates ca. 1 UV lesion/nucleosome (200 bp) in 15% of the nuclear volume. Irradiated cells were allowed to recover in culture medium for 2 h before fixation (in the presence of 50 µM 5-aza-2’-deoxycytidine for DNMT immunodetection).

### UVC laser irradiation

Cells were grown on quartz coverslips (25 mm diameter, thickness No.1, Nevco) and nuclei were stained by adding Hoechst 33258 (10 μg/ml final, Sigma-Aldrich) to the culture medium 45 min before UVC irradiation. Quartz coverslips were transferred to a Chamlide magnetic chamber (Gataca-systems) on a custom stage insert (Live Cell Instrument) and cells were irradiated for 40 ms using a 2 mW pulsed diode-pumped solid-state laser emitting at 266 nm (RappOptoElectronics, Hamburg GmbH) directly connected to a Zeiss LSM900 confocal microscope adapted for UVC transmission with all-quartz optics in a 37°C and 5% CO_2_ incubation chamber. The laser was attenuated using a 0.1% neutral density filter and focused through a 40x/0.6 Ultrafluar glycerol objective with quartz lenses. The laser is controlled by a UGA Firefly module with SysCon2 software (Rapp OptoElectronic). The laser impact has an average size of 1 μm in diameter and damages around 1% of the total nuclear volume. The corresponding UVC dose, not directly measurable but estimated by comparing the intensity of the CPD damage generated by the laser and the UVC lamp, is 500 J/m^2^.

### Labelling S-phase cells with EdU

Cells grown on glass coverslips (Thorlabs) were incubated in culture medium with EdU (5-ethynyl-2’-deoxyuridine, Life Technologies) at a final concentration of 10 μM for 15 minutes at 37°C followed by three washes in PBS. Cells were further kept in medium without EdU until fixation with 2% paraformaldehyde (Euromedex) for 12 minutes at room temperature. The EdU was detected using the Click-it EdU Alexa Fluor 647 imaging kit (Life technologies) according to manufacturer’s instructions.

### Labelling repair synthesis with EdU

Cells grown on glass coverslips (Thorlabs) and irradiated with the UVC lamp through a micropore filter, were incubated with EdU (5-ethynyl-2’-deoxyuridine, Life Technologies) at a final concentration of 10 μM for 2 h at 37°C, followed by three washes in PBS. Cells were then fixed with 2% paraformaldehyde (Euromedex) for 12 minutes at room temperature. The EdU was detected using the Click-it EdU Alexa Fluor 647 imaging kit (Life Technologies) according to manufacturer’s instructions. S-phase cells with pan-nuclear EdU signal were excluded from the analysis.

### Cell proliferation assays

HCT-116-DNMT1-AID2 and HCT-116-UHRF1-AID2 cells were seeded in 6-well plates at 5x10^3^cells/ml and 1x10^4^ cells/ml, respectively. Two days later, cells were treated or not with 1 μM auxin analogue 5-Ph-IAA (referred to as auxin for simplicity) and exposed to global UVC irradiation (10J/m^2^) 2 hours later in PBS containing auxin or not. A non-irradiated control was also included. Cells were left to grow in culture medium with or without auxin for 20 hours, before auxin wash out. Viable cells were counted with an automated cell counter (Fluidlab R-300 cell counter, Anvajo) 3, 6 and 8 days after seeding.

### Micronuclei scoring

U2OS cells were plated on glass coverslips at a concentration of 8x10^4^ cells/ml one day before global UVC irradiation with a UVC lamp at 10 J/m^2^. The DNMT1 inhibitor GSK-3484862 was added to the culture medium 3 h before UV irradiation and kept 2 h after. Cells were then left to recover in fresh medium for 30 h before fixation with 2% paraformaldehyde (Euromedex) for 12 minutes, permeabilisation with 0.2% Triton X-100 (Euromedex) in PBS for 5 min, and mounting of the coverslips in Vectashield medium with DAPI (4’,6-diamidino-2-phenylindole, Vector laboratories). Micronuclei were scored with a Leica DMI6000 epifluorescence microscope.

### Immunofluorescence

Cells grown on coverslips were fixed with 2% paraformaldehyde (Euromedex) for 12 minutes at room temperature. In most cases, cells were pre-extracted before fixation with 0.5% Triton X-100 in CSK buffer (Cytoskeletal buffer: 10 mM PIPES pH 7.0, 100 mM NaCl, 300 mM sucrose, 3 mM MgCl_2_) for 5 min at room temperature. When no pre-extraction was done, fixed cells were permeabilized with 0.2% Triton X-100 (Euromedex) in PBS for 5 min at room temperature. To detect 5mC or 5hmC, DNA was denatured with HCl (WWR chemicals) as indicated in **Supplementary Table S3**. HCl was neutralized with 4 washes in PBS.

Samples were blocked for 10 min in 5% BSA (Bovine Serum Albumin, Sigma-Aldrich) in PBT (PBS 0.5% Tween-20), and incubated for 45 min at room temperature with the appropriate primary antibodies (**Supplementary Table S4**) diluted in blocking buffer. After three washes in PBT, samples were incubated for 30 min at room temperature with secondary antibodies coupled to AlexaFluor 488, 568, 594, or 647 (**Supplementary Table S4**) diluted in blocking buffer. Finally, coverslips were mounted in Vectashield medium or Vectashield plus antifade medium with DAPI (4’,6-diamidino-2-phenylindole, Vector laboratories).

For immunofluorescence combined with EdU detection, the EdU Click-it reaction was performed either before (U2OS cells) or during immunofluorescence, between the primary and secondary antibodies because it affects the GFP signal (NIH/3T3 GFP-DDB2 cells). GFP immunodetection was used to enhance GFP signal. The EdU Click-it reaction was systematically followed by a second fixation step and permeabilization.

### SNAP-tag labelling of histones

For specific labelling of newly synthesized histones cells were grown on glass coverslips and pre-existing SNAP-tagged histones were first quenched by incubating cells with 10 μM of the non-fluorescent substrate SNAP-cell Block (New England Biolabs) for 30 min followed by a 30 min-wash in fresh medium and a 2 h chase. The new SNAP-tagged histones synthesized during the chase were fluorescently labelled with 2 μM of the red-fluorescent reagent SNAP-cell TMR star (New England Biolabs) during a 15 min-pulse step followed by 45 min wash in fresh medium. Cells were subsequently permeabilised with Triton X-100, fixed and processed for immunostaining. Cells were irradiated with a UVC lamp before the pulse step.

### Cell extracts and western blot

Total extracts were obtained by scraping cells on plates in Laemmli buffer (50 mM Tris-HCl pH 6.8, 1.6% SDS (Sodium Dodecyl Sulfate), 8% glycerol, 4% β-mercaptoethanol, 0.0025% bromophenol blue) followed by 5 min denaturation at 95°C. Total protein concentration was measured with a NanoDrop (DeNovix), and then the extracts were diluted accordingly to load equal protein amounts.

For western blot analysis, extracts were run on 4%–20% Mini-PROTEAN TGX gels (Bio-Rad) in running buffer (200 mM glycine, 25 mM Tris, 0.1% SDS) and transferred onto nitrocellulose membranes (0.2 μm, Amersham Protran) for 30 min at 15V with a Trans-Blot SD semidry transfer cell (Bio-Rad). Total proteins were revealed by Pierce® Reversible Stain (Thermo Scientific). Proteins of interest were probed using the appropriate primary and HRP (Horse Radish Peroxidase)-conjugated secondary antibodies (**Supplementary Table S4**), and detected using SuperSignal West Pico or Femto chemiluminescence substrates (Pierce). The resulting signal was visualized with the Odyssey Fc imaging system (LI-COR Biosciences).

### Isolation of genomic DNA for Nanopore sequencing

Cells were seeded in 15-cm dishes to harvest at least 5 million cells per condition on the day of the experiment. Replication was inhibited with 10 mM Hydroxyurea (HU, Sigma-Aldrich) and 1 mM cytosine arabinoside (Ara-C, Sigma-Aldrich) in fresh growth medium for 2 h at 37°C prior to global UVC irradiation (10 J/m^2^) in PBS supplemented with 10 mM HU and 1 mM Ara-C to maintain the inhibition of replication. A non-irradiated sample was used as negative control. Cells were trypsinized and counted and cell pellets were washed once in PBS. DNA isolation from cell pellets was performed with the Monarch HMW DNA Extraction Kit for Cells & Blood (New England Biolabs) following manufacturer’s instructions. Genomic DNA was then fragmented to achieve an average size of 7-9 kb with g-TUBEs (Covaris) following manufacturer’s instructions.

### Nanopore sequencing and data analysis

DNA library preparation from genomic DNA was done using the Native Barcoding Kit 24 V14 (SQK-NBD114-24, Oxford Nanopore Technologies), starting from 2 μg DNA for each sample. Multiplexing was performed to sequence 4 samples at the same time. The samples were run on the Promethion sequencing device (Oxford Nanopore Technologies), with flow cell type FLO-PRO114M and Software MinKNOW version 24.11.10. Adaptive sampling was performed focusing on the genomic regions listed in **Supplementary Table S5** (hs1 coordinates).

Base calling was performed with the Dorado base calling program (dorado-0.9.5 Oxford Nanopore Technologies), using the model dna_r10.4.1_e8.2_400bps_sup@v5.0.0_5mCG_5hmCG@v3, filtering with min-qscore 10 and mapping on hg38. DNA methylation was analyzed with the Methylator program (https://methylator.readthedocs.io/en/latest/) developed by the in-house BiBS bioinformatics platform. Configuration parameters are provided with the raw data files.

### IPOND-R (Isolation of Proteins On Nascent DNA at Repair sites)

#### EdU labelling of repair sites

Cells were seeded in 15-cm dishes to harvest at least 70 million cells per condition on the day of the experiment. In order to label only repair patches, replication was inhibited with 10 mM Hydroxyurea (HU, Sigma-Aldrich) and 1 mM cytosine arabinoside (Ara-C, Sigma-Aldrich) in fresh growth medium for 2 h at 37°C prior to global UVC irradiation (10 J/m^2^) in PBS supplemented with 10 mM HU and 1 mM Ara-C to maintain the inhibition of replication. A non-irradiated sample was used as negative control. After irradiation, cells were incubated in fresh medium containing the replication inhibitor cocktail and 10 µM Ethynyl-deoxyUridine (EdU, Euromedex) for 1 h at 37°C.

#### Cell fixation and permeabilization

Cells were fixed with 1% Formaldehyde (Sigma-Aldrich) for 15 minutes on a rocking platform. The fixation was stopped by adding 1.33 M Glycine (Sigma-Aldrich) for 5 minutes on the rocking platform. Cells were washed twice in cold PBS, scraped in 1% BSA solution in PBS and pelleted for 5 min at 300 g at 4°C. Cell pellets were permeabilized with 1% Triton X-100 (1 ml/ 10 million cells, Euromedex) for 30 min at room temperature and washed twice with cold 1% BSA solution.

#### EdU biotinylation by Click-it chemistry

Cell pellets were resuspended in a Click-it reaction cocktail containing 2 mM CuSO_4_ (Sigma-Aldrich), 1 mM Tris((1-hydroxy-propyl-1H-1,2,3-triazol-4-yl)methyl)amine (THPTA, Euromedex), 10 mM Sodium Ascorbate (Sigma-Aldrich), 5 mM Amino-guanidine hydrochloride (Sigma-Aldrich), and 10 μM Biotin Picolyl azide (Sigma-Aldrich). Cell suspensions were left on a rotating wheel at room temperature for 1 h. Cells were then washed twice in cold 1% BSA solution containing 0.5X EDTA-free Protease inhibitor cocktail (PIC, Roche), and once in cold PBS.

#### Cell lysis and sonication

Cell pellets were resuspended in lysis buffer (200 µl per 15 million cells) containing 1% SDS, 50 mM Tris pH 7.5, and 1X PIC. The cell suspensions were transferred to 1.5 ml sonication tubes (Diagenode) and sonication was performed using a Bioruptor Pico (Diagenode) with 4-8 cycles of 30 sec ON and 30 sec OFF. The sonicated samples were transferred to new Eppendorf tubes and centrifuged 10 min at 16,000 g at 4°C. The supernatants were transferred to Low Protein Binding tubes (Thermo Fisher Scientific).

#### DNA isolation and analysis of DNA shearing

A 20 μl aliquot of the -UV and +UV samples was incubated with 200 mM NaCl and 200 μg/ml RNAse A (Millipore) overnight at 65°C. After addition of 100 μg/ml Proteinase K (Sigma-Aldrich), samples were incubated for a further 2 h at 45°C prior to DNA purification using a PCR purification kit (Macherey-Nagel) following manufacturer’s instructions. DNA samples were analyzed with a TapeStation (Agilent) to check that the average fragment size was about 500 bp.

#### Streptavidin capture of biotin-labelled nascent DNA and associated proteins

Each sample was resuspended in an equal volume of cold PBS supplemented with 1 X PIC to dilute the SDS present in the sonication buffer. An input sample (0.1%) was taken at this stage. For streptavidin capture, Dynabeads MyOne Streptavidin-C1 beads (Invitrogen) were washed in lysis buffer and in PBS supplemented with 1X PIC and then added to sonicated cell lysates (1 µl beads per 1 million cells) and incubated overnight at 4°C on a rotating wheel. After capture, beads were washed once in lysis buffer (1 ml wash solution per 10 million cells), once in 1M NaCl supplemented with 1X PIC, followed by 3 additional washes in lysis buffer.

#### Protein analysis in the capture

The beads were resuspended in Laemmli buffer. Laemmli buffer was also added to the input samples and decrosslinking was achieved by heating for 5 min at 95°C. Samples were analyzed by western blot.

### RNA extraction and mRNA sequencing

#### RNA extraction

U2OS or HCT116 DNMT1-AID2 cells were seeded in 6-well plates to reach 10^6^ cells per well on the day of RNA extraction. RNA extraction was performed on cell pellets using RNAeasy MiniKit (Qiagen 74014) following manufacturer’s instructions.

#### mRNA sequencing

RNA sequencing was performed at the Genomi’IC Platform of Institut Cochin (Paris, FR). Total RNA (around 500 ng) was used for library preparation. Only RNA samples with an RNA Integrity Number (RIN) greater than 7 were used to ensure high-quality input material. Library preparation was performed using either the Illumina Stranded mRNA Prep Ligation Kit or the NEBNext Ultra II Directional RNA Library Prep Kit (New England Biolabs), following manufacturer’s instructions. In short, polyadenylated mRNA molecules were selectively enriched using oligo(dT) magnetic beads. The captured mRNA was then reverse transcribed into complementary DNA (cDNA), preserving the strand orientation of the original transcripts. Subsequently, Illumina-compatible adapters were ligated to the cDNA fragments, followed by PCR amplification to generate the final libraries.

#### mRNA sequencing analysis

RNA sequencing analysis was performed utilizing RASflow_EDC v1.3 (https://github.com/parisepigenetics/RASflow_EDC), adapted from RASflow [50]. In short, quality control was performed and the reads were trimmed to remove low quality sequences and adaptors. HISAT2 [51] was used for alignment on hg38. Gene counts were performed with featureCounts (with -M --fraction options) [52] using GeneCode annotation (v41). DESeq2 Package [53] was exploited to run differential expression analysis. Principal Component Analysis was done using the regularized log count data. Genes were considered significantly differentially expressed with an adjusted p-value <0.05. The complete list of parameters and tool versions is available in the raw data files. The workflow was run on the HPC cluster of iPOP-UP, hosted by Ressource Parisienne en Bioinformatique Structurale (RPBS). Gene ontology analysis was performed with enrichR (https://maayanlab.cloud/Enrichr/) using the MSigDB_Hallmark_2020 database [54–56].

### Image acquisition and analysis

Fluorescence imaging was performed with a Leica DMI6000 epifluorescence microscope using a PlanApochromat 40x/1.3 or 63x/1.4 oil objective. Images were captured using a CCD camera (Teledyne Photometrics) and Metamorph software, and mounted using Adobe Photoshop. Live cell imaging coupled to UVC laser micro-irradiation was performed using a 40x/0.6 Ultrafluar Glycerol objective on a Zeiss LSM900 confocal microscope. Images were captured using Zen blue software, and analysed with ImageJ (U. S. National Institutes of Health, Bethesda, Maryland, USA, http://imagej.nih.gov/ij/). The subtract background function was applied to all images prior to quantification, and images were smoothed using the median filter function. Nuclei and heterochromatin domains were segmented based on DAPI staining, and UVC-damaged regions based on GFP-DDB2 fluorescence or immunostaining for UV damage or repair factors using the default threshold function with manual adjustment. Mean intensities were measured in each region of interest using the measure and set measurement functions. The enrichment or depletion of proteins at UV sites were quantified by dividing the mean intensity in the UV-damaged area of the nucleus by the mean intensity in the undamaged part of the nucleus (total nucleus subtracted from damaged area).

### Statistical Analysis

Volcano plots were generated with ggplot2 on RStudio 2023.03.1. Statistical tests were performed using GraphPad Prism. Enrichments/depletions at UV sites were transformed in a log2 base and were compared to a theoretical mean of 0 in one sample t-tests. Comparisons between two groups were quantified using Student’s t-tests with Welch’s correction. Multiple comparisons were quantified using one-way or two-way ANOVA with Dunnett’s or Tukey’s post-test or using Kruskal-Wallis test with Dunn’s post-test in case of non-gaussian distribution of the values. Student’s t-tests and ANOVA analyses were run on the means of independent experiments. Comparisons of proliferation curves were based on non-linear regression with a second order polynomial model. ns: non-significant, *: p < 0.05, **: p < 0.01, ***: p < 0.001, ****: p < 0.0001.

## RESULTS

### Maintenance of DNA methylation marks at sites of UV damage repair

As a first approach to determining whether any gross alterations of DNA methylation would arise after UV damage repair, we performed DNA sequencing with the nanopore technology to decipher 5mC and 5hmC changes at the single base level upon UV damage induction in human cells (**Fig. 1A**). U2OS cells were globally irradiated with a UVC lamp at 10 J/m^2^, which generates UV lesions broadly distributed through the genome, every 6 kb on average [57]. DNA damage induction by UV irradiation was controlled through increased γH2A.X levels in western blot (**Fig. 1A**). We used nanopore sequencing with adaptive sampling for higher coverage (up to 26X), focusing on 3% of the genome including highly methylated regions such as HOX gene clusters and rDNA repeats. We detected 5mC in around 60% of the sequenced CpGs in control non-irradiated cells and did not reveal any significant change in average 5mC levels 1 h and 4 h post UV irradiation (**Fig. 1B**, upper panels). 5mC analysis was also performed by dividing the genome into 500 bp tiles with a minimum coverage of 10X, of different 5mC density, from fully methylated (100% methylation, all CpGs harbour a methyl group) to unmethylated (0% methylation, no CpG harbors a methyl group). Comparing the non-irradiated control with 1 h and 4 h post UV samples, no significant difference was detected in 5mC distribution. 5hmC levels were also analyzed in a similar manner. As expected [58], very low levels of 5hmC were detected, and we did not find any significant alteration in 5hmC amounts or distribution after UV irradiation (**Fig. 1B**, lower panels).

**Figure 1.**
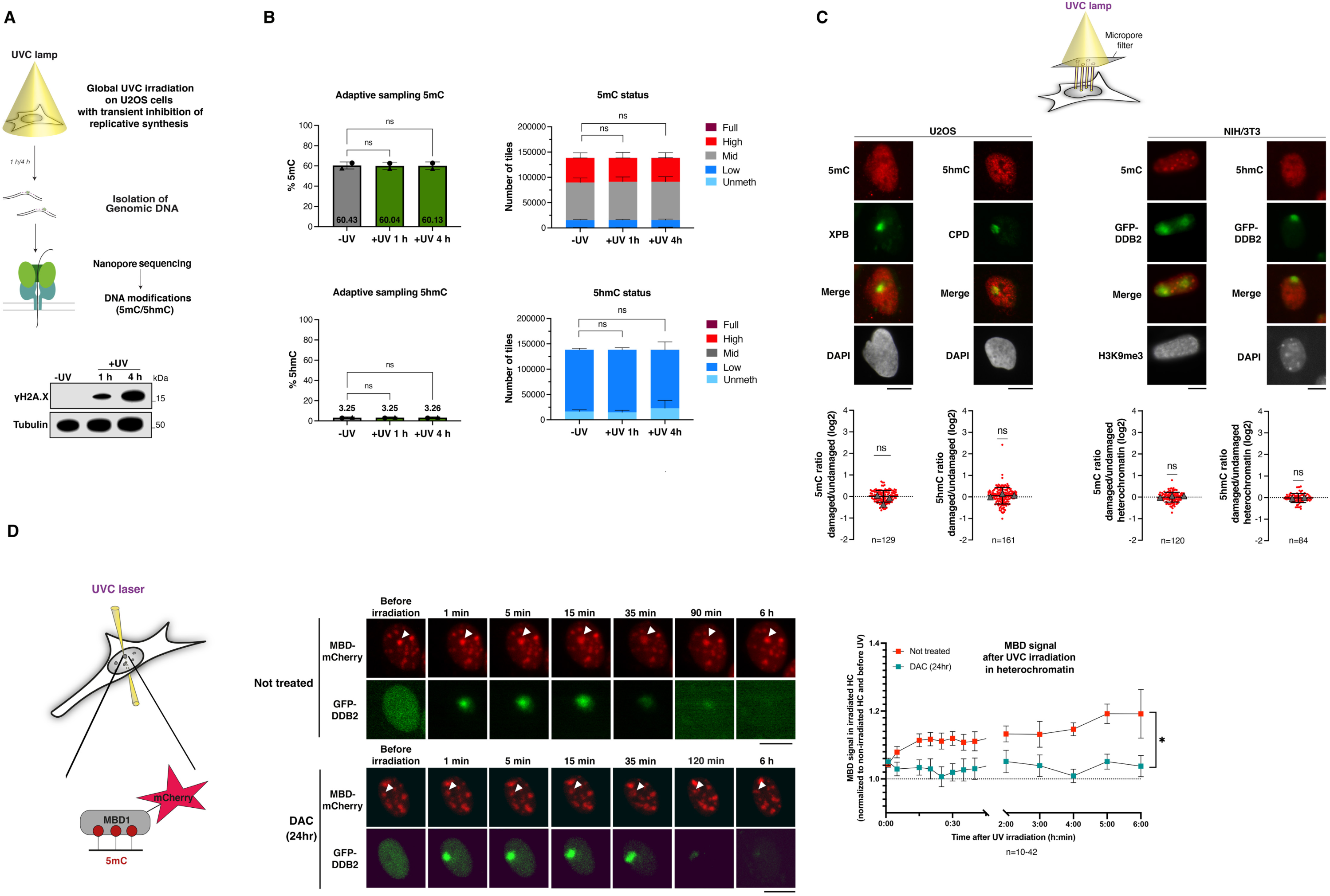
Maintenance of DNA methylation marks at sites of UV damage repair in mammalian cells. **(A)** Scheme of the methodology used to isolate UV-damaged DNA for subsequent Nanopore sequencing of DNA methylation marks (5mC/5hmC). UV damage induction is controlled by western blot for the DNA damage marker γH2A.X (Tubulin, loading control). **(B)** Left: 5mC (upper panel) and 5hmC (lower panel) average levels in genomic regions of interest (adaptive sampling) in cells exposed to UV irradiation (+UV) compared to non-irradiated cells (-UV) are shown in percentages. Right: DNA methylation status by sample is shown in number of tiles (Full: 100% methylated; High: 75%-100% methylated; Mid 25%-75% methylated; Low 0-25% methylated; Unmeth:100% unmethylated). Tile size: 500 bp. Results from two independent experiments. **(C)** Quantification of 5mC and 5hmC at UV damage sites (marked with CPD, XPB or GFP-DDB2) analyzed by immunofluorescence 2 h after local UVC irradiation in human U2OS cells and in mouse NIH/3T3 GFP-DDB2 cells. In NIH/3T3 GFP-DDB2 cells, H3K9me3 staining is combined with 5mC staining to recognize heterochromatin domains that are not visible with DAPI staining due to the harsh HCl pre-treatment. The ratio of 5mC or 5hmC mean intensities in the damaged vs undamaged regions of the same nucleus is transformed in a log2 scale so that positive values correspond to enrichment, and negative values to depletion at UV sites. Data are presented as mean values ± SD from n cells scored in two or three representative experiments. Triangles represent the mean values of each independent experiment. Data are presented in a log2 scale so that positive values correspond to enrichment, and negative values to depletion at UV sites. **(D)** DNA methylation at UV damage sites monitored by live cell imaging in NIH/3T3 cells expressing GFP-DDB2 (damage sensor) and MBD-mCherry (5mC reader) and treated or not with decitabine (DAC) 24h before UV irradiation. The white arrowhead indicates the damaged heterochromatin (HC) domain. MBD level quantification is performed by measuring the integrated density of the MBD signal in the damaged heterochromatin domain normalized to an undamaged heterochromatin domain in the same nucleus and relative to before UV irradiation. Data are presented as mean values from n cells ± SEM. Statistical analysis is performed by one-way or two-way ANOVA (B), one-sample Student’s t-test compared to a hypothetical value of 0 (C) or through the comparisons of areas under the curves (D). Scale bars, 10 μm.

As subtle alterations of DNA methylation marks within a short repair patch of 26-27 nucleotides in length [59] may be too diluted to be detected by genome-wide sequencing, we decided to focus on local changes specifically at UV damage sites through imaging experiments in cells subject to localized damage infliction. First, we exploited antibodies against 5mC and 5hmC in immunofluorescence in mammalian cells exposed to local UVC irradiation through micropore filters (**Fig. 1C**). 5mC and 5hmC immunodetection required DNA denaturation, achieved through HCl treatment, and 5mC and 5hmC signal specificity was verified after 48 h treatment with the DNMT inhibitor 5-aza-2’-deoxycytidine (decitabine, DAC), which also activates the TET demethylation pathway, leading to the expected decrease of 5mC and increase of 5hmC immunofluorescence signals (**Supplementary Fig. S1A**) [60,61]. 5mC and 5hmC staining was performed 2 h after local UVC irradiation of the cells, i.e. when damage excision and repair synthesis are initiated, with co-staining for UV photoproducts (cyclobutane pyrimidine dimers, CPD) or for the UV damage repair factor XPB (Xeroderma Pigmentosum, complementation group B) to mark sites of UV damage in human U2OS cells. Quantification of the 5mC and 5hmC signal at UV damage sites relative to the rest of the nucleus did not reveal any significant enrichment or depletion of DNA methylation marks at UV sites (**Fig. 1C**, left panel). We next focused our analysis on DNA methylation-enriched heterochromatin domains by conducting similar experiments in mouse cells, where pericentric heterochromatin domains are visualized by DAPI or H3K9me3 staining. For this, we employed NIH/3T3 fibroblasts engineered to express a GFP-tagged UV damage sensor DDB2 (DNA damage binding protein 2), which marks UV damage sites [46]. Focusing on UV lesions overlapping with heterochromatin domains, we compared 5mC and 5hmC levels in damaged vs. undamaged heterochromatin. Even in the heterochromatin context, neither 5mC nor 5hmC showed any significant depletion or accumulation at UV damage sites (**Fig. 1C**, right panel). To further analyze DNA methylation dynamics at UV sites in native conditions, we established an antibody-free approach for real-time tracking of DNA methylation in live cells. For this, we transfected NIH/3T3 GFP-DDB2 mouse cells with a fluorescently tagged 5mC reader, the methyl-binding domain (MBD) of MBD1 (mCherry-tagged construct adapted from [62]). MBD-mCherry showed the expected enrichment in highly methylated pericentric heterochromatin, which was erased upon 48 h treatment with the DNMT inhibitor decitabine (DAC), validating this tool for monitoring alterations in 5mC levels (**Supplementary Fig. S1B**). Exploiting a confocal microscope coupled with a UVC laser, we performed targeted micro-irradiation of pericentric heterochromatin, visualized by Hoechst staining in live nuclei. GFP-DDB2 accumulation and release was used as a proxy for UV damage induction and repair, and we followed the dynamics of MBD-enriched heterochromatin domains over time after UVC laser damage. Damaged heterochromatin domains decompacted quickly after UV damage and recompacted during repair progression, as previously reported [46] (**Supplementary Fig. S1C**). To determine whether the decompaction of damaged heterochromatin domains was accompanied by a change in 5mC levels, we quantified the MBD signal in UV-damaged heterochromatin, normalized to a non-irradiated area in the same nucleus. We did not detect any loss of MBD signal in irradiated heterochromatin domains but rather a modest increase (**Fig. 1D**). To assess whether this change in MBD levels was determined by DNMT activity, we treated cells for 24 hours with decitabine (DAC). This short inhibition of DNMTs did not result in a loss of DNA methylation in heterochromatin domains **(Supplementary Fig. S1D)** but prevented the MBD level increase following UVC laser micro-irradiation in heterochromatin even though heterochromatin decompaction was comparable to untreated cells (**Supplementary Fig. S1C**), arguing that DNMTs are involved in DNA methylation control in UV-damaged heterochromatin.

Overall, these analyses reveal a maintenance and/or fast re-establishment of DNA methylation marks during UV damage repair in mammalian cells, possibly accompanied by de novo methylation events.

### Recruitment of the DNA methylation machinery to UV damage sites

To uncover the mechanisms that control DNA methylation during DNA damage repair, we examined the recruitment of the DNA methylation machinery to repair sites. For this, we explored a proteomic dataset that we previously obtained by mass spectrometry analysis of captured DNA repair patches in UV-damaged U2OS cells [63]. This revealed an enrichment at sites of UV damage repair of several factors known to be involved in DNA methylation maintenance at replication forks: the DNA methyltransferase DNMT1, its co-factor UHRF1 and their interactors PCNA, LIG1 and PAF15 (**Fig. 2A**, volcano plot). We confirmed the presence of DNMT1 and UHRF1 at repair patches by western blot analysis of captured proteins and also identified the de novo methyltransferase DNMT3A by these means (**Fig. 2A**, western blot).

**Figure 2.**
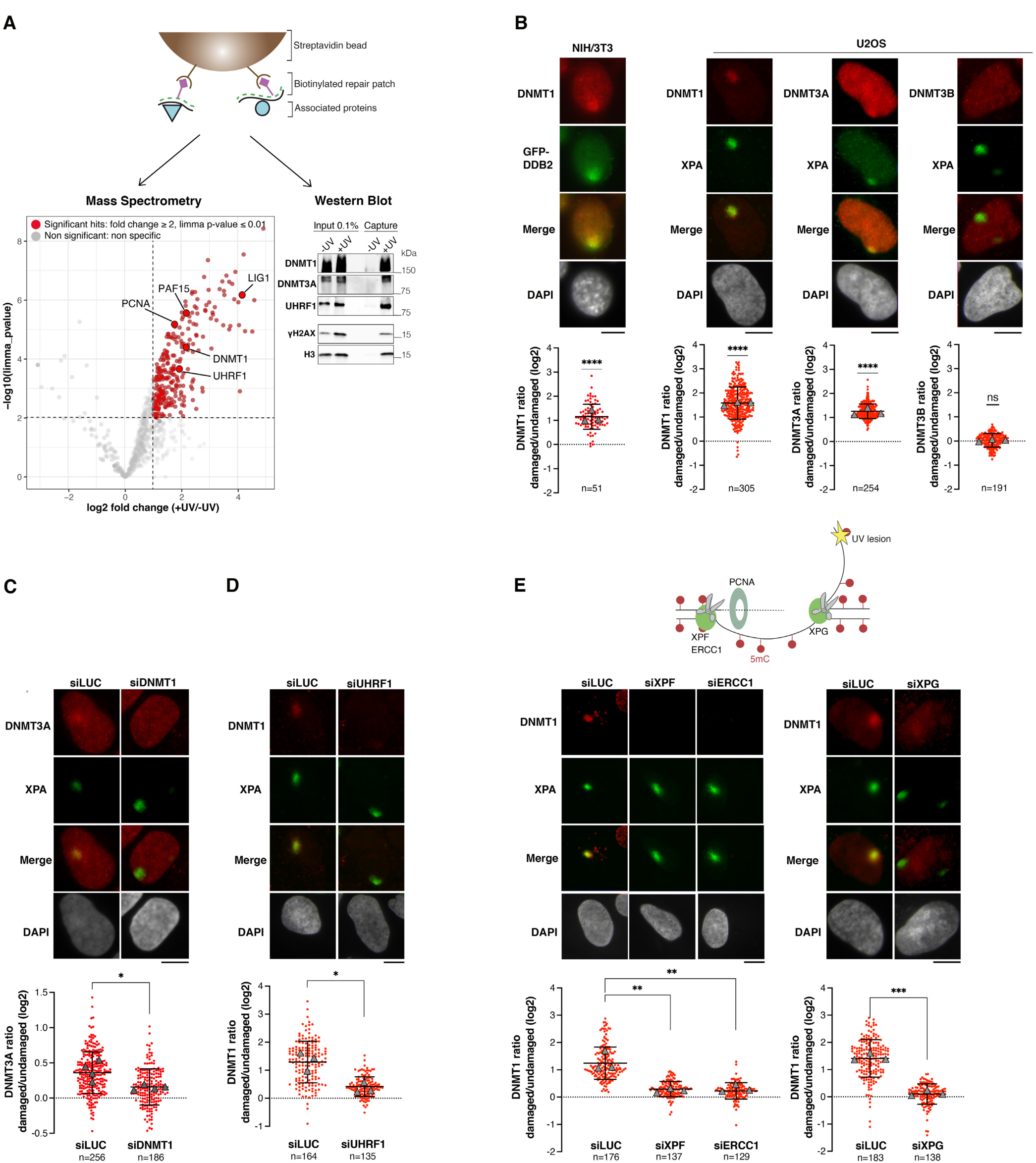
Recruitment of the DNA methyltransferases DNMT1 and DNMT3A to sites of UV damage repair. **(A)** Volcano plot showing the proteome of UV damage repair patches analyzed by mass spectrometry 1 h after UVC irradiation in U2OS cells. Proteins significantly enriched in irradiated compared to non-irradiated samples (+/-UV fold-change ≥ 2 and limma p-value ≤ 0.01) are shown in red. Factors known to be involved in DNA methylation dynamics are highlighted and their presence in the +UV capture is validated by western blot. **(B)** Recruitment of DNMTs (red) to UV damage sites (green) analyzed by immunofluorescence 2 h after local irradiation with a UVC lamp in mouse NIH/3T3 GFP-DDB2 cells and human U2OS cells. S-phase cells showing a focal pattern of DNMTs (and replication-associated EdU incorporation, not shown) were excluded from the analysis. **(C-E)** DNMT3A and DNMT1 recruitment to UV damage sites (marked by XPA) analyzed by immunofluorescence 2 h after local UVC irradiation in U2OS cells treated with the indicated siRNAs (siLUC, control). The scheme in panel (E) shows the nucleases XPF-ERCC1 and XPG involved in UV damage excision. Scale bars, 10 μm. Graphs show mean values ± SD from n cells scored in 3 independent experiments. Triangles represent the mean values of each independent experiment. Data are presented in a log2 scale so that positive values correspond to enrichment, and negative values to depletion at UV sites. Statistical analysis is performed with limma (A), one-sample Student’s t-test compared to a hypothetical value of 0 (B), two-sided Student’s t-test with Welch’s correction (C-E) or one-way ANOVA (E).

We further investigated the recruitment of DNMTs to UV damage sites by immunofluorescence in cells exposed to local UVC irradiation using specific antibodies to detect DNMTs (**Supplementary Fig. S2A**) while UV sites were marked by proteins involved in UV damage recognition and repair. During the 2 h recovery post irradiation, cells were treated with the cytidine analogue decitabine (DAC), which captures active DNMTs on DNA thus facilitating the visualization of their enrichment at UV sites (**Supplementary Fig. S2B**). DNMT1 showed a pronounced accumulation at repair sites both in mouse and human cells, while DNMT3A enrichment was more modest and DNMT3B recruitment was not detectable (**Fig. 2B**). Since the accumulation of DNMT1 and DNMT3A at UV sites was observed in decitabine-treated cells and in conditions where soluble proteins were removed by detergent extraction prior to cell fixation, these DNMTs were present in an active chromatin-bound form at sites of UV damage repair. We next dissected their order of recruitment through siRNA-mediated loss-of-function approaches (**Supplementary Fig. S2C**), which showed that DNMT1 was recruited independently of DNMT3A, but was necessary for DNMT3A recruitment (**Fig. 2C** and **Supplementary Fig. S2D**). UHRF1 allows DNMT1 recruitment to hemi-methylated DNA during replicative synthesis [5]. We therefore tested whether UHRF1 was similarly required for DNMT1 accumulation at repair sites. Corroborating this hypothesis, UHRF1 depletion abrogated DNMT1 enrichment at sites of UV damage repair (**Fig. 2D**) with no detectable effect on DNMT1 total levels as measured by western blot (**Supplementary Fig. S2E**).

To further characterize the molecular mechanisms controlling DNMT1 recruitment to UV sites, we investigated a possible dependency on repair synthesis during the late steps of the NER pathway. To do so, we targeted the endonucleases XPF-ERCC1 and XPG, involved in the excision of the damaged oligonucleotide prior to repair synthesis [64]. The knock-down of these proteins was confirmed by western blot on total cell extracts and did not measurably affect DNMT1 total levels but abrogated repair synthesis at UV sites as expected (**Supplementary Fig. S2F-G**). DNMT1 recruitment to UV damage sites was significantly decreased by knock-down of XPF-ERCC1 and XPG (**Fig. 2E**). These results demonstrate that DNMT1 is recruited to UV damage sites following damage excision.

UHRF1 enrichment at UV sites was tested with specific antibodies (**Supplementary Fig. S2H**) (**Fig. 3A**). In contrast to DNMT1, UHRF1 recruitment at UV sites was not dependent on the presence of XPF-ERCC1 and XPG (**Fig. 3B**). To dissect how UHRF1 was recruited to UV sites, we expressed GFP-tagged UHRF1 constructs [11] in U2OS cells (single UHRF1 domains or the full-length protein as a control). Only the Tandem Tudor domain (TTD) and, to a lesser extent, the Plant homeodomain (PHD) of UHRF1, showed a significant enrichment at UV damage sites as observed with the full-length protein (**Fig. 3C**), indicating that both domains are sufficient for UHRF1 recruitment. TTD and PHD are known to interact with di- and tri-methylated H3 on lysine 9, and di- and tri-methylated LIG1 on lysine 126, and the methyltransferase responsible for depositing H3K9me2 and LIG1K126me2 is G9a/GLP [11]. Consistent with the role of the TTD and PHD domains in UHRF1 recruitment, cells treated with a G9a/GLP inhibitor showed a reduced recruitment of UHRF1 at UV sites (**Fig. 3D**, left panel). G9a/GLP inhibition reduced H3K9 methylation levels, as expected, but did not impact UHRF1 total levels in the nucleus (**Supplementary Fig. S2I).**

**Figure 3.**
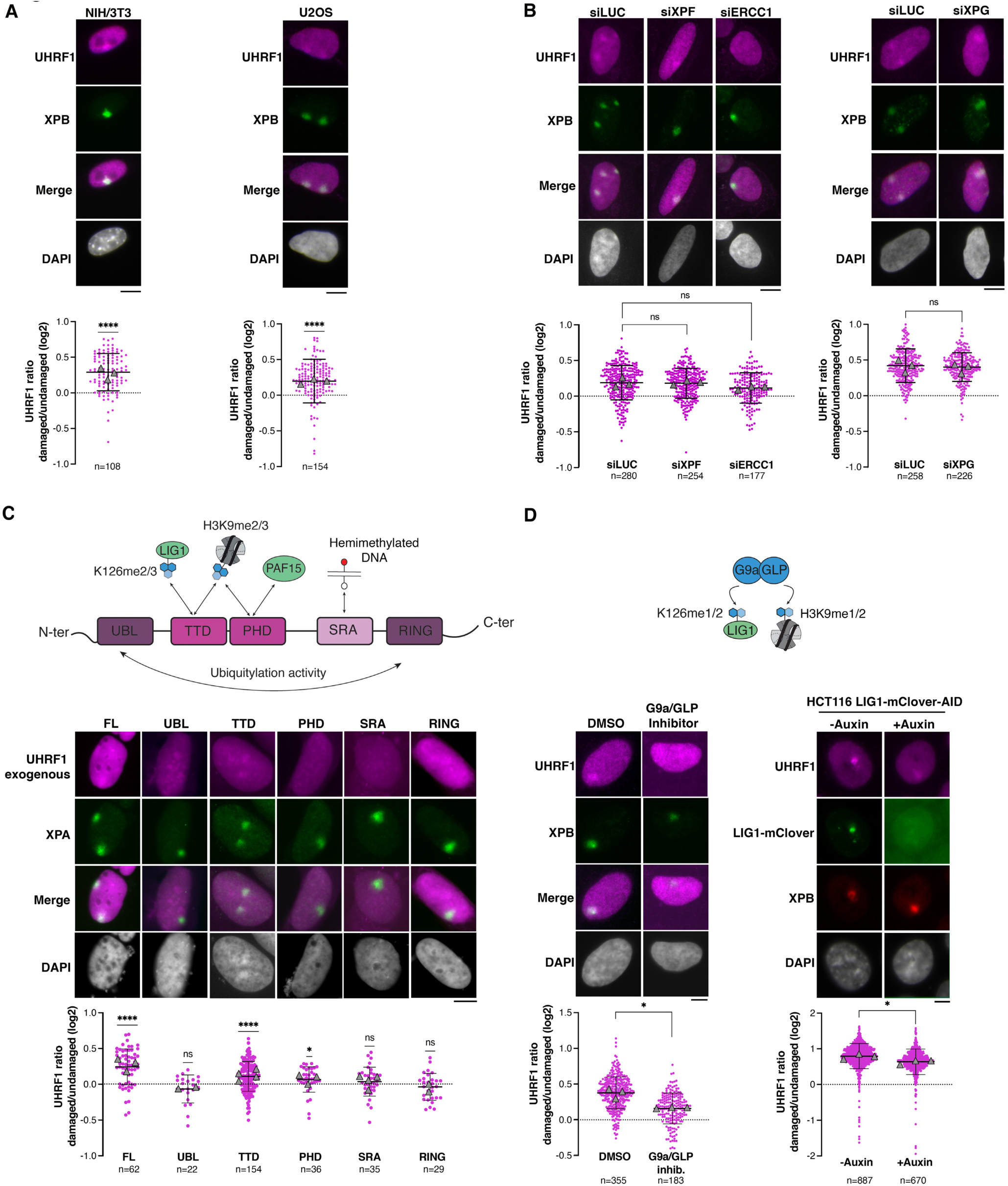
Recruitment of the DNMT1 co-factor UHRF1 to sites of UV damage repair. **(A)** Recruitment of UHRF1 (purple) to UV damage sites (green) analyzed by immunofluorescence 2 h after local irradiation with a UVC lamp in mouse NIH/3T3 GFP-DDB2 cells and human U2OS cells. **(B)** UHRF1 recruitment to UV sites (marked by XPB) analyzed by immunofluorescence 2 h after local UVC irradiation in U2OS cells treated with the indicated siRNAs (siLUC control). **(C)** Recruitment of fluorescent exogenous UHRF1 (full length, FL or single domains) to UV sites (marked by XPA) analyzed as in (A) in U2OS cells transiently expressing UHRF1 constructs. The scheme represents UHRF1 domains and their known interactors. **(D)** Left panel: UHRF1 localization at UV sites (marked by XPB) analyzed as in (A) after 72 h treatment with a G9a/GLP inhibitor. The scheme represents known targets of G9a/GLP methyl-transferases. Right panel: UHRF1 localization at UV sites (marked by XPB) analyzed by immunofluorescence 2 h after local irradiation with a UVC-laser coupled to a confocal microscope in HCT116 LIG1-mClover-AID2 cells treated or not with auxin. Scale bars, 10 μm. Graphs show mean values ± SD from n cells scored in 2 to 4 independent experiments. Triangles represent the mean values of each independent experiment. Data are presented in a log2 scale so that positive values correspond to enrichment, and negative values to depletion at UV sites. Statistical analysis is performed by one-sample Student’s t-test compared to a hypothetical value of 0 (A, C), two-sided Student’s t-test with Welch’s correction (B, D) or one-way ANOVA (B).

To determine which G9a/GLP target among LIG1 and histone H3 was necessary for UHRF1 recruitment at UV sites, we exploited an HCT116 cell line engineered with a LIG1-mClover-auxin-inducible degron (AID2) system, which allows a complete degradation of LIG1 upon a short treatment with the auxin analogue 5-Ph-IAA without decreasing UHRF1 total levels **(Fig. 3D, Supplementary Fig. S2J**). LIG1 depletion led to a modest but significant decrease in UHRF1 enrichment at UV sites, hinting towards a minor contribution of LIG1 to UHRF1 recruitment at UV damage sites (**Fig. 3D**, right panel).

Together, these findings demonstrate that the DNA methylation machinery is recruited to sites of UV damage repair with an accumulation of both maintenance and de novo DNMTs, dependent on the DNMT1 co-factor UHRF1 and on UV damage excision.

### Recruitment of the DNA methylation machinery depends on histone deposition at sites of UV damage repair

UV damage repair is accompanied by chromatin rearrangements, which include the deposition of newly synthesized histones by specific histone chaperones [65]. In particular, the histone chaperone complexes CAF-1 and HIRA deposit new H3.1 and H3.3 histone variants, respectively, at sites of UV damage repair [66,67]. We thus wondered whether histone dynamics at UV damage sites may influence the recruitment of the DNA methylation machinery and reciprocally. To test the dependency of the DNA methylation machinery on newly deposited H3, we individually knocked down the chaperones CAF-1 and HIRA, or H3.3 directly by targeting its two gene products. HIRA and H3.3 knockdowns did not impact DNMT1 and UHRF1 total levels (**Supplementary Fig. S3A**). CAF-1 depletion in contrast led to a significant reduction in UHRF1 total levels (**Supplementary Fig. S3A**) and H3.1 could not be directly depleted as it is encoded by 10 genes in mammalian cells. We thus focused our analysis on HIRA and H3.3 depletions and tested their impact on the DNA methylation machinery at UV sites. Both HIRA and H3.3 knockdowns strongly decreased DNMT1 recruitment to UV sites as assessed by immunofluorescence (**Fig. 4A**, left panel).

**Figure 4.**
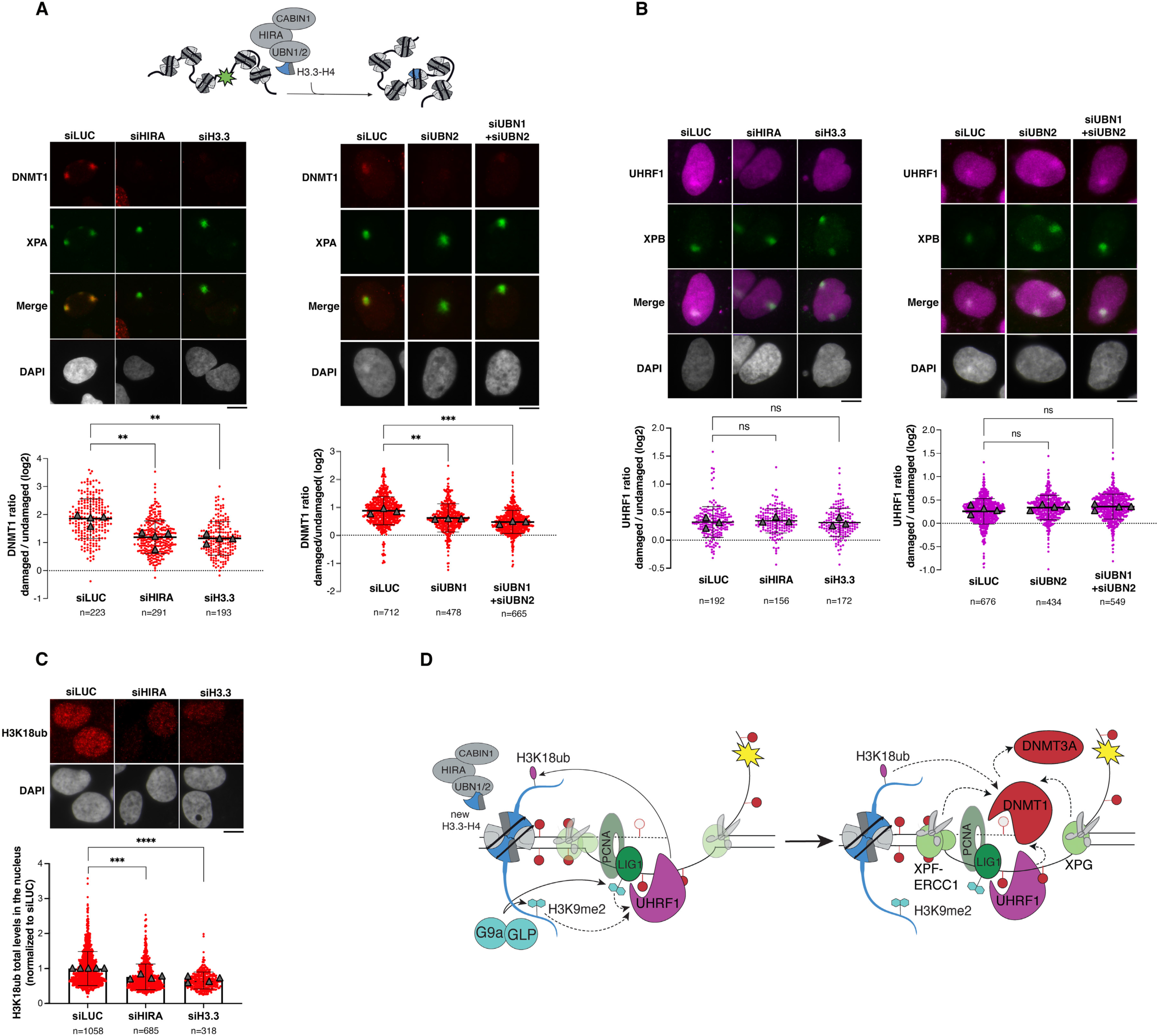
Recruitment of the DNA methylation machinery depends on new histone deposition at sites of UV damage repair. **(A)** Recruitment of DNMT1 (red) to UV damage sites (green) analyzed by immunofluorescence 2 h after local UVC irradiation in U2OS cells treated with the indicated siRNAs (siLuc, control). The scheme shows new H3.3 deposition by the histone chaperone complex HIRA in UV-damaged chromatin. **(B)** Recruitment of UHRF1 (purple) to UV damage sites (green) analyzed by immunofluorescence 2 h after local UVC irradiation in U2OS cells treated with the indicated siRNAs (siLuc, control). **(C)** H3K18ub levels (red) analyzed by immunofluorescence in U2OS cells treated with the indicated siRNAs (siLuc, control). **(D)** Scheme illustrating the hierarchical recruitment of the DNA methylation machinery to sites of UV damage repair and the crosstalks with DNA repair factors and with histone modifications and variants. Plain arrows indicate modifying activities and dotted arrows indicate protein recruitment. Scale bars, 10 μm. Graphs show mean values ± SD from n cells scored in 3 to 4 independent experiments. Triangles represent the mean values of each independent experiment. In (A) and (B), data are presented in a log2 scale so that positive values correspond to enrichment, and negative values to depletion at UV sites. In (C), data are presented as mean values in the nucleus normalized to the control (siLUC). Statistical analysis is performed by one-way ANOVA.

HIRA is a complex composed of three core subunits, CABIN1, HIRA, and UBN1 or UBN2 histone chaperones, which are necessary for new H3.3 deposition while old H3.3 recycling instead depends on HIRA binding to the ASF1a/b histone chaperone [68,69]. To directly test the effect of new H3.3 deposition on DNMT1 recruitment to UV sites, we thus depleted UBN1/2 alone or in combination (**Supplementary Fig. S3A-B**). While UBN1 knockdown did not significantly affect DNMT1 recruitment to UV sites (**Supplementary Fig. S3C**), UBN2 depletion impaired DNMT1 recruitment and the effect was even more pronounced in condition of double UBN1 and UBN2 knockdowns (**Fig. 4A**, right panel). In all cases, total levels of DNMT1 and UHRF1 remained unchanged (**Supplementary Fig. S3A**). These results point to a specific dependency of DNMT1 recruitment on de novo H3.3 deposition at UV damage sites. UHRF1 enrichment at UV damage sites was not impacted when we interfered with the de novo deposition of H3.3 (**Fig. 4B**) but total chromatin-bound levels of UHRF1 in the nucleus were reduced in siHIRA and siH3.3 conditions (**Fig. 4B, Supplementary Fig. S3D**).

To further decipher how HIRA-mediated deposition of newly synthesized H3 variants at UV sites might drive DNMT1 recruitment, we focused on H3 modifications that are recognized by DNMT1. We analyzed the histone mark H3K18ub, which is deposited by UHRF1, and read by DNMT1 at replication foci [70]. We first verified the specificity of H3K18ub immunofluorescence signal, which showed an increase upon DNMT1 depletion in a UHRF1-dependent manner (**Supplementary Fig. S3E**) as reported [71]. Next, we observed that H3K18ub nuclear levels were significantly decreased by HIRA and H3.3 depletions (**Fig. 4C**). Together, these observations suggest that HIRA-mediated deposition of new H3.3 promotes DNMT1 recruitment to UV damage sites by enhancing UHRF1 binding to chromatin thus driving H3K18ub that is recognized by DNMT1.

While HIRA-mediated deposition of H3.3 promoted DNMT1 recruitment to UV damage sites, DNMT1 had no reciprocal impact on HIRA accumulation at those sites (**Supplementary Fig. S3F**). Knockdown of the DNA methylation machinery by targeting DNMT1 or UHRF1 also did not affect new H3.3 deposition at UV sites as monitored through SNAP-tag based labelling of new histones (**Supplementary Fig. S3G**).

Collectively, these findings reveal multiple layers of control for the recruitment of the DNA methylation machinery to sites of UV damage repair, which relies on deposition of new H3 histones by the HIRA chaperone, on specific H3 post-translational modifications by G9a/GLP and UHRF1, and on UV damage excision by the repair machinery (**Fig. 4D**).

### Functional relevance of maintaining DNA methylation upon UV damage

Knowing the importance of DNA methylation in controlling gene expression and cell identity, we set out to decipher the functional relevance of DNA methylation maintenance during UV damage repair.

First, we observed that a functional DNA methylation machinery was not required for UV damage repair, as DNMT1 and UHRF1 depletions did not inhibit repair synthesis, which we monitored by EdU incorporation at repair sites in U2OS cells (**Supplementary Fig. S4A**). Consistent with this finding, genome integrity was not impaired by transient inhibition of DNMT1 around the time of UV irradiation (using the non-nucleoside DNMT1-specific inhibitor GSK-3484862, DNMT1i). We scored the formation of micronuclei as a proxy for genomic instability. While UV irradiation increased the proportion of micro-nucleated cells as expected, DNMT1 inhibition did not exacerbate this effect (**Supplementary Fig. S4B**).

We then examined whether a loss-of-function of the DNA methylation machinery through DNMT1 or UHRF1 degradation could sensitize the cells to UV irradiation. We performed proliferation assays in HCT116 cell lines engineered with a DNMT1- or UHRF1-auxin-inducible degron (AID2) system, which allows complete degradation of DNMT1 or UHRF1 upon a short treatment with the auxin analogue 5-Ph-IAA (referred to as auxin for simplicity) 2 h prior to UVC irradiation (**Fig. 5A**). Auxin was washed out 20 h after irradiation to allow the majority of UV lesions to be repaired in the absence of a functional DNA methylation machinery. While auxin treatment had a modest inhibitory effect on cell growth in non-irradiated cells, the inhibition of cell proliferation was exacerbated in UV-damaged cells (**Supplementary Fig. S4C, Fig. 5B**), arguing that a functional DNA methylation machinery during UV damage repair is crucial for cell proliferation.

**Figure 5.**
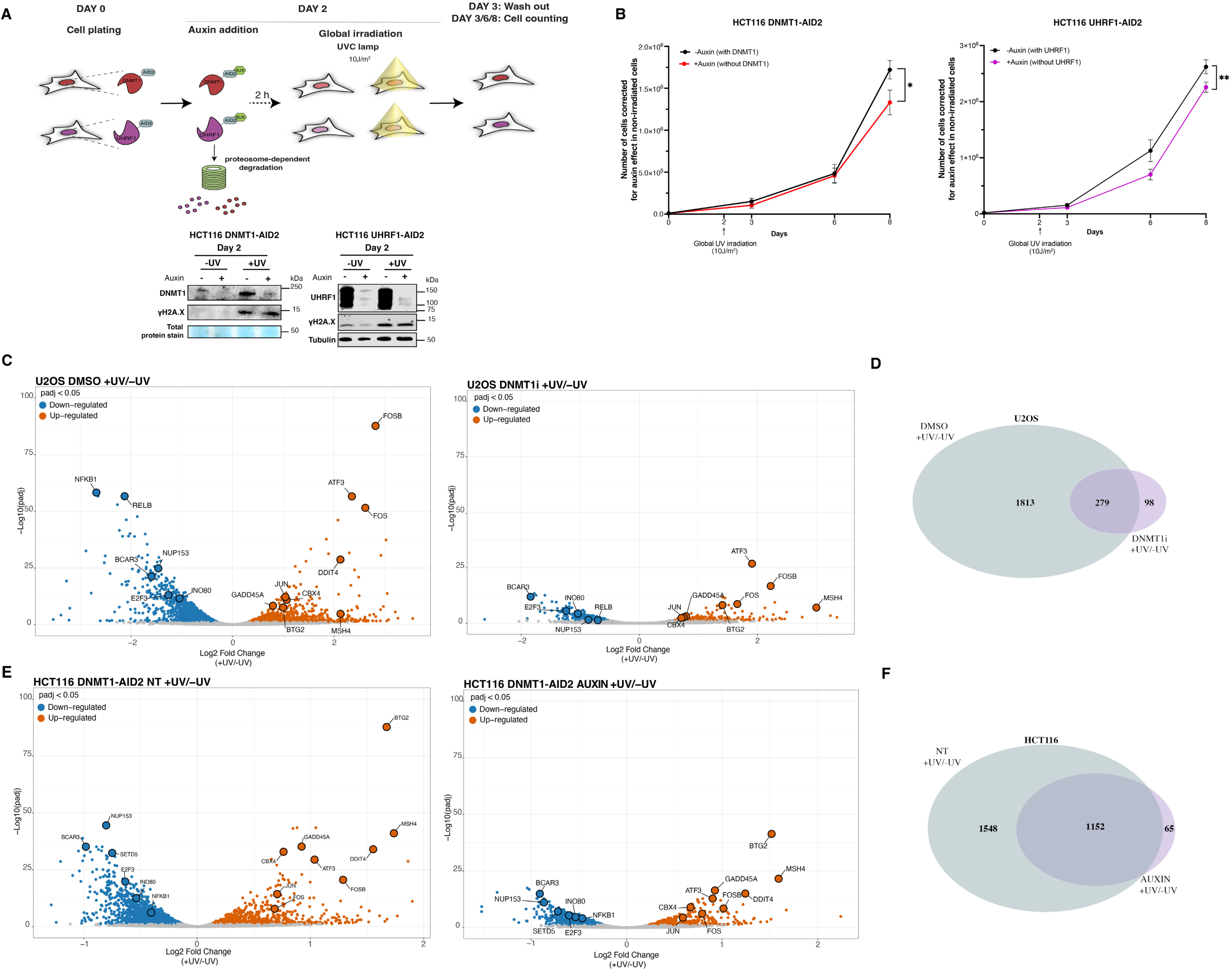
Functional relevance of DNA methylation maintenance following UV damage. (A) Experimental scheme of proliferation assays post UV irradiation in HCT116-DNMT1-AID2 and HCT116-UHRF1-AID2 cells. The Auxin-induced degradation of DNMT1 and UHRF1 and DNA damage induction by UV irradiation (monitored by γH2A.X) are controlled by western blot at day 2 (total protein stain and Tubulin are used as loading controls). **(B)** Proliferation assays in HCT116-DNMT1-AID2 (left) and HCT116-UHRF1-AID2 cells (right) exposed to Auxin treatment and global UVC irradiation (10 J/m^2^) as shown on the experimental scheme. The graphs show mean values ± SEM from 4 (DNMT1) and 5 (UHRF1) independent experiments. The number of cells in the +Auxin condition is corrected for the effect of Auxin in non-irradiated cells (shown in Supplementary Fig. S4C). Statistical analysis is performed by non-linear regression with a second order polynomial model. **(C)** Volcano plots showing the transcriptome changes 2 h post global UVC irradiation in U2OS cells in control conditions (DMSO) and conditions where DNMT1 was inhibited around the time of damage exposure (DNMT1i). Genes significantly up- or down-regulated in irradiated compared to non-irradiated samples (+/-UV adjusted p-value <0.05) are shown in red or blue, respectively. Some factors known to be differentially expressed upon UV irradiation are annotated. **(D)** Venn diagram representing the number of Differentially Expressed Genes (DEG) between irradiated and non-irradiated U2OS cells (+UV/-UV) in control (DMSO, grey) and DNMT1 inhibitor (DNMT1i, violet) conditions. DEG are considered significant if the adjusted p-value is <0.05. **(E)** Volcano plots showing the transcriptome changes 2 h post global UVC irradiation in HCT116-DNMT1-AID2 cells in control conditions (NT) and conditions where DNMT1 was inhibited around the time of damage exposure with auxin (AUXIN). Genes significantly enriched or depleted in irradiated compared to non-irradiated samples (+/-UV adjusted p-value <0.05) are shown in red or blue, respectively. Some factors known to be differentially expressed upon UV irradiation are highlighted. **(F)** Venn diagram representing the number of Differentially Expressed Genes (DEG) between irradiated and non-irradiated HCT116-DNMT1-AID2 cells (+UV/-UV) in control (NT, grey) Auxin treated cells (AUXIN, violet) conditions. DEG are considered significant if the adjusted p-value is <0.05.

To further dissect the effect of DNMT1 inhibition on the UV-damage response, we analyzed the transcriptome of UV-damaged cells (2 hours post UV irradiation) compared to control cells. Intriguingly, short-term treatment of cells with the catalytic inhibitor of DNMT1 around the time of UV irradiation markedly affected the transcriptional response to UV irradiation as revealed by mRNA sequencing in U2OS cells (**Fig. 5C**). Experiments were performed in triplicates and reproducibility was controlled by principal component analysis (**Supplementary Fig. S4D**). In control cells (DMSO-treated), we detected the expected upregulation of immediate early genes known to be early transcribed upon UV irradiation in mammalian cells, such as ATF3, JUN, FOS and BTG2 (**Fig. 5C**, left panel) [72–74]. We also detected the upregulation of known stress-responsive genes such as DDIT4 and GADD45A [75,76], and among down-regulated genes, we detected expected hits such as the cell cycle regulator E2F3, or inflammatory genes RELB and NFKB1 [74]. Gene ontology analysis of differentially expressed genes (DEG) upon UV irradiation showed an enrichment of signaling pathways known to respond to UV damage in mammalian cells such as MAPK and TNF pathways (**Supplementary Fig. S4E**, left panel) supporting the validity of our transcriptome analysis [77,78]. Similar pathways were enriched in the analysis of cells treated with a DNMT1 inhibitor around the time of UV irradiation (**Supplementary Fig. S4E** right panel), but the amplitude and significance/reproducibility of the gene expression changes and thus the total number of differentially expressed genes were much lower (**Fig. 5C-D**). Such short-term DNMT1 inhibition did not trigger major transcriptional changes in non-irradiated cells (**Supplementary Fig. S4F**) but it led to a dampened and unscheduled transcriptional response to UV damage. We confirmed these results by mRNA sequencing in HCT116 DNMT1-AID2 cells, which were treated with auxin 2 hours before irradiation, to achieve complete DNMT1 degradation at the time of DNA damage induction (**Supplementary Fig. S4G-H**). Auxin was kept until 2 hours post irradiation, when cells were harvested for mRNA analysis. In control cells which express DNMT1, the response to UV damage induction was comparable to that of U2OS cells treated with DMSO, with up and down-regulation of expected genes and enrichment of similar signaling pathways as revealed by gene ontology analysis (**Fig. 5E**, **Supplementary Fig. S4I**). Upon DNMT1 degradation by auxin treatment, the amplitude and significance of UV-induced gene expression changes was reduced as was the number of differentially expressed genes (**Fig. 5E-F**), confirming a dampened and unscheduled transcriptional response to UV damage in the absence of a functional DNA methylation machinery. DNMT1 degradation on its own did not result in significant transcriptional changes in non-irradiated cells (**Supplementary Fig. S4J**).

Together, these analyses reveal the importance of an active DNA methylation maintenance machinery for the transcriptional response to UV damage and for sustained cell proliferation.

## DISCUSSION

DNA methylation plays a central role in controlling gene expression and genome integrity. Stable inheritance of DNA methylation through somatic cell divisions is thus of major importance to maintain cell identity. Here, by dissecting the fundamental mechanisms controlling DNA methylation dynamics following UV damage in mammalian cells, we have deciphered a machinery recruited at UV damage sites to ensure DNA methylation maintenance during UV damage repair (**Fig. 4D**). We reveal the hierarchical recruitment of UHRF1, DNMT1 and DNMT3A and their functional crosstalk with UV damage repair, and with the surrounding chromatin dynamics, including new H3 histone deposition and post-translational modifications.

### Machinery for DNA methylation maintenance at UV sites

By employing three complementary methodologies to analyze DNA methylation marks, we did not detect any loss of 5mC upon UV irradiation, but rather a modest increase in irradiated heterochromatin. We can therefore speculate that during UV damage repair, the majority of 5mC methylation marks are rapidly and faithfully re-established on the newly synthesized strand through the concerted action of UHRF1, DNMT1 and DNMT3A. DNMT1 presumably copies pre-existing 5mC onto the newly synthesized strand at repair sites. The modest increase in DNA methylation that we detect at UV sites could be explained by some de novo DNA methylation by DNMT3A, or by the presence of a 3-strand DNA repair intermediate including the newly synthesized repaired strand and the not yet excised damaged oligonucleotide. DNMT3A might also cooperate with DNMT1 in DNA methylation maintenance at UV sites, as reported at specific loci in mouse embryonic stem cells and post-implantation embryos [79,80] and at DNA double-strand breaks in human cells [24].

The DNA methylation machinery that we identify at UV sites is similar to the one acting at replication forks to re-establish DNA methylation marks on newly synthesized DNA [5] with some specificities: DNMT3A is recruited at repair sites but does not have a canonical role during replication; while during DNA replication LIG1 methylation plays a crucial role for UHRF1 recruitment [11], UHRF1 recruitment to UV sites is only modestly impacted by LIG1 depletion and thus appears to be mostly reliant on the other G9a/GLP target, which is H3K9; UHRF1 is recruited to UV damage sites via its TTD domain, which is dispensable for DNA methylation maintenance in a replicative context [81].

While DNMT1 recruitment to UV sites depends on UV damage excision, UHRF1 recruitment to sites of UV damage repair is not mediated by XPF-ERCC1 even though an interaction was reported with this nuclease complex [32]. UHRF1 roles in the DNA damage response were so far confined to interstrand crosslink and DNA double-strand break repair and involved its SRA and RING domains [28,31,32]. Here, we demonstrate that UHRF1 is also involved in the UV damage response, through its TTD domain.

Besides 5mC maintenance by DNMTs, TET oxidative enzymes may also contribute to regulating DNA methylation marks at UV sites. While we do not detect major changes in 5hmC levels, TET enzymes were shown to mediate a 5hmC increase in response to different types of DNA damage inducers, among which UV rays [33,34,45]. Moreover, in the absence of exogenous damage, the NER protein XPC has been shown to promote 5mC to 5hmC transition [45]. This hints towards a possible connection of UV damage repair proteins with active DNA demethylation, which will be worth exploring in future studies.

### Crosstalk with histone dynamics at UV sites

We have deciphered a crosstalk between new H3.3 histone deposition at UV sites and the DNA methylation machinery, showing that new histone deposition promotes DNMT1 recruitment to UV sites. These intriguing findings suggest that DNA methylation maintenance coordinates with new histone dynamics at sites of UV damage repair. Newly synthesized histones bearing distinct post-translational modifications from old histones[82], they may be more prone to ubiquitylation by UHRF1, thus favoring DNMT1 recruitment. In addition, we have observed that newly deposited H3.3 histones promote UHRF1 binding to chromatin, which increases H3K18ub, further stimulating DNMT1 recruitment. We cannot exclude that a similar mechanism might also be driven by newly deposited H3.1 histones, but interfering with H3.1 deposition impacted the cell cycle and UHRF1 levels too much to be able to dissect UV-specific mechanisms.

Further investigation of other H3 ubiquitination marks deposited by UHRF1 could be envisioned to thoroughly analyze the crosstalk between new H3 deposition and H3 ubiquitination driving DNMT1 recruitment.

Besides repair-coupled chromatin dynamics, it is conceivable that the chromatin environment of the damaged locus may govern the mechanisms and fidelity of DNA methylation maintenance, with possible differences between highly methylated heterochromatin and poorly methylated euchromatin.

### Functional relevance of DNA methylation maintenance at UV sites

DNMT1 and UHRF1 degradation both impede the proliferation of UV-damaged cells. Consistent with our data, UHRF1-depleted cells were shown to be more sensitive to several types of DNA damage, including UV rays [28,30,32,83]. Together, these findings highlight the importance of a functional DNA methylation machinery during UV damage repair for cell growth/survival. Upon DNMT1 inhibition in U2OS cells or degradation in HCT116 DNMT1-AID2 cells, we also detect an unscheduled and dampened transcriptional response to UV damage. This may result from DNA methylation alterations in damaged genes in the absence of the DNA methylation maintenance machinery. It may also relate to other functions of DNMT1, independent of DNA methylation maintenance at UV damage sites. Indeed, DNMT1 was recently shown to control mRNA modifications and stability [84], and we cannot exclude that DNMT1 performs a similar post-transcriptional role post UV damage.

Further strengthening the functional role of DNA methylation maintenance coupled to repair, Xeroderma Pigmentosum (XP) patients, who do not have a functional NER machinery, also show a generally hypomethylated genome [85]. The dependency of DNMT1 recruitment to UV sites on UV damage excision by the NER machinery could be an explanation for the hypomethylation observed in XP genomes. DNA misrepair would thus be a source of DNA methylation alterations that could promote pathological phenotypes. Whether this change in DNA methylation is directly linked to defective DNA methylation re-establishment during UV damage repair, or arises from subsequent genomic instability is still unknown.

Overall, our study puts forward a network of factors involved in maintaining DNA methylation at sites of DNA damage, thus illuminating how DNA damage repair can protect from DNA methylation alterations like those associated with diseases. Altered DNA methylation re-establishment upon chronic/repetitive DNA damage might indeed trigger persistent changes in DNA methylation that impact gene expression, cell identity, and disease onset.

## ACKNOWLEDGEMENTS

We thank members of our laboratory for stimulating discussions and Masato Kanemaki for providing the HCT116-LIG1-AID2 cell line. We acknowledge the help of the Functional Epigenomics EpiG platform for access to instruments and technical advice, the Bioinformatics and Biostatistics Core Facility BiBs for bioinformatic analyses and the Imaging Platform Epi^2^ for confocal and epifluorescence microscopy (Epigenetics and Cell Fate Center).

## AUTHOR CONTRIBUTIONS

M.M, S.P., L.G., S.E.P designed and performed experiments and analyzed the data. M.M. and S.E.P. wrote the manuscript. M.M., L.F., M.F., M.H., performed the Nanopore sequencing experiments. M.H. and M.F. analyzed the Nanopore sequencing data using bioinformatic tools established by O.K and E.B. M.M. and M.H. analyzed the RNA sequencing data. L.F. generated UHRF1 constructs and K.Y. generated DNMT1 and UHRF1 degron cell lines in P-A.D. laboratory.

## CONFLICT OF INTEREST

The authors declare no competing interests.

## FUNDING

Work in S.E.P. laboratory was supported by the European Research Council (consolidator grant ERC-2018-CoG-818625 “REMIND”), the “Who am I?” laboratory of excellence (ANR-11-LABX-0071) funded by the French Government through its “Investments for the Future” program (ANR-11-IDEX-0005-01), the French National Research Agency (ANR-24-CE12-3079-04) and the Fondation pour la Recherche Medicale (EQU202503020067). Work in P.A.D. laboratory was supported by the Agence Nationale de la Recherche (ANR-23-CE12-0015-01), the Institut National du Cancer (INCA_18350), and the Fondation pour la Recherche Medicale (EQU202503019988).

The bioinformatics analyses were performed on the HPC cluster of iPOP-UP, hosted by RPBS and funded by the Université Paris Cité (IDEX). M.M. received PhD and postdoctoral fellowships from University Paris Diderot, La Ligue Contre le Cancer and EUR G.E.N.E (ANR-17-EURE-0013).

## DATA AVAILABILITY

Nanopore sequencing data and RNA sequencing data is available from the European Nucleotide Archive (ENA) with accession number PRJEB121327. Other raw data files are deposited in the Figshare repository: https://doi.org/10.6084/m9.figshare.33018800

## Notes

### Competing Interest Statement

The authors have declared no competing interest.

