## Supplementary figures and tables for "New histone deposition recruits the DNA methylation maintenance machinery at sites of DNA damage repair"

#### Supplementary Figure S1

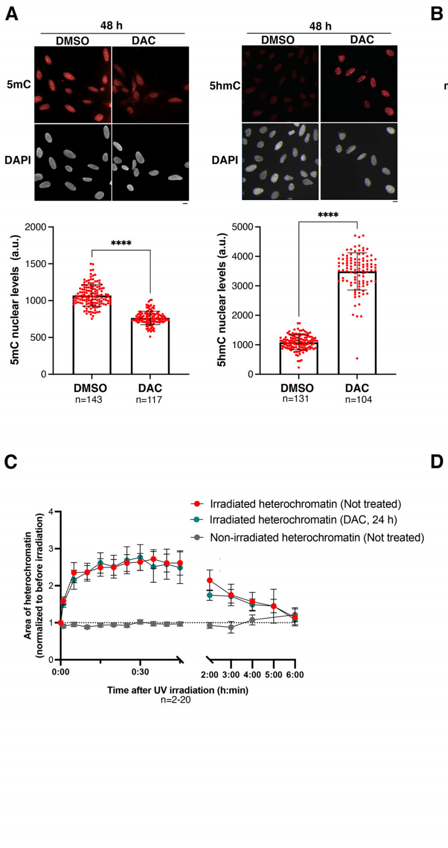

**Supplementary Figure S1: Validation of 5mC and 5hmC detection tools**

**(A)** Quantification of 5mC and 5hmC analyzed by immunofluorescence in U2OS cells treated for 48 h with decitabine (DAC) or with a control (DMSO).

(**B**) The specific detection of DNA methylation with the MBD-mCherry construct is validated with 48 h decitabine treatment (DAC), which erases the enrichment in heterochromatin domains.

(**C**) The decompaction of UV-irradiated heterochromatin is quantified by measuring the area of the heterochromatin domain along time relative to before irradiation. The area of a non-irradiated heterochromatin domain is shown as a control.

(**D**) Control that MBD-mCherry enrichment in heterochromatin domains is preserved upon 24 h decitabine (DAC) treatment.

Scale bars, 10 µm. Data on the graphs are presented as mean values ± SD from n cells or ± SEM from n cells in (C). Statistical analysis is performed with a two-sided Student t-test. a.u., arbitrary units

### Supplementary Figure S2

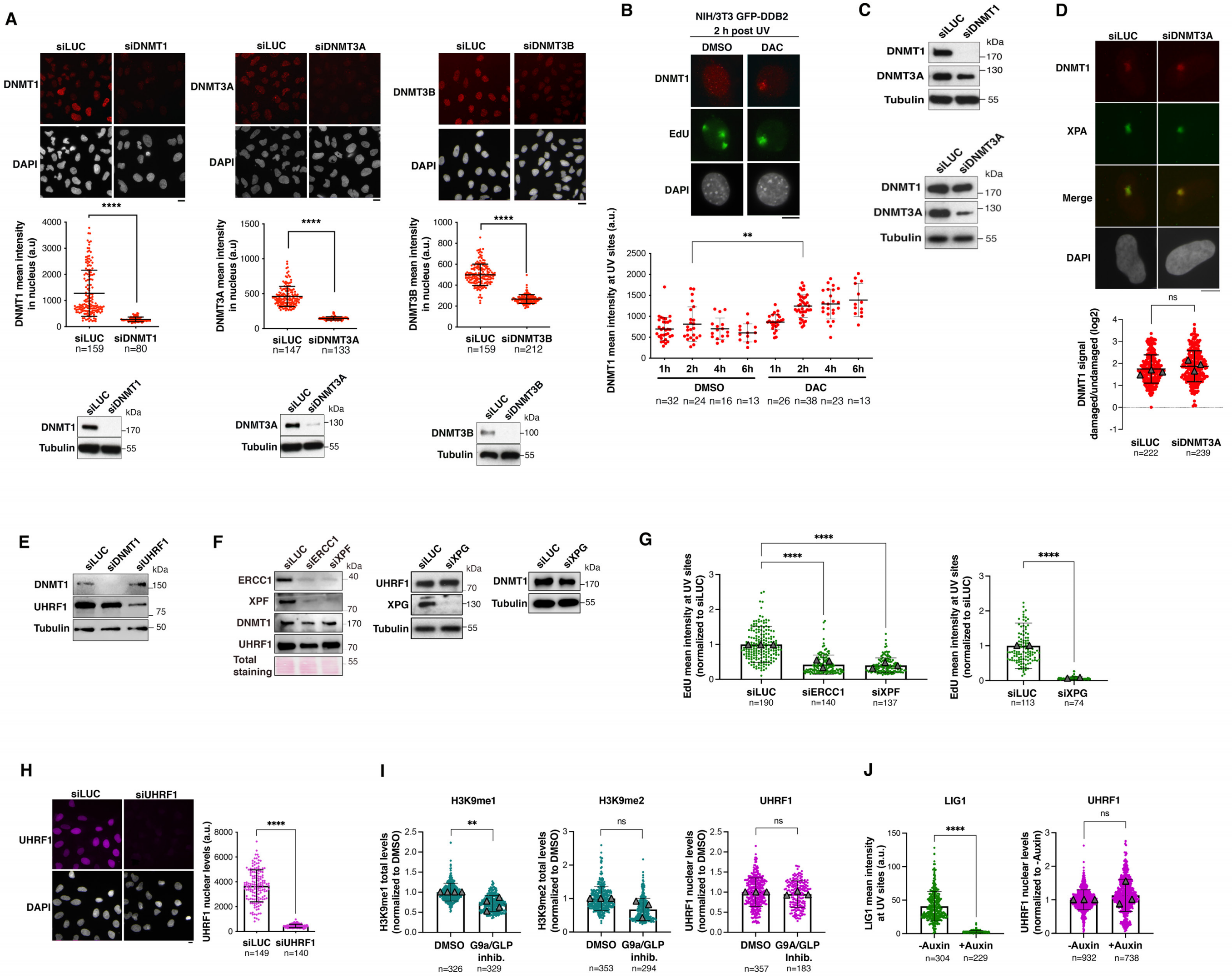

**Supplementary Figure S2: Analysis of the DNA methylation machinery at sites of UV damage repair.**

(**A**) Validation of DNMT1, DNMT3A, and DNMT3B antibodies by immunofluorescence and western blot in U2OS cells treated with the respective siRNAs (siLUC, control). Western blot is performed on total cell extracts (Tubulin, loading control).

(**B**) Quantification of DNMT1 mean intensity at UV sites (marked by EdU) in NIH/3T3 GFP-DDB2 cells treated with decitabine (DAC) or DMSO (vehicle) for the indicated times after UV irradiation.

**(C, E, F)** Western blot on total cell extracts from U2OS cells treated with the indicated siRNAs (siLUC, control). Tubulin or total protein staining are shown as loading controls.

**(D)** Recruitment of DNMT1 (red) to UV damage sites (green) analyzed by immunofluorescence 2 h after local UVC irradiation in U2OS cells treated with siDNMT3A (siLUC, control).

**(G)** Quantification of EdU levels at sites of local UVC irradiation in U2OS cells treated with the indicated siRNAs (siLUC, control).

**(H)** Validation of UHRF1 antibody by immunofluorescence in U2OS cells treated with the indicated siRNAs (siLUC, control).

(**I**) Quantification of H3K9me1, H3K9me2, and UHRF1 nuclear levels in U2OS cells treated with the G9a/GLP inhibitor UNC0642 (1 μM for 72 h; DMSO, control).

(**J**) Quantification of LIG1 levels at UV sites in HCT116 LIG1-mClover-AID cells treated or not with Auxin (1 μM for 2 h) and exposed to UVC laser micro-irradiation.

Scale bars, 10 μm. Data on the graphs are presented as mean values ± SD from n cells. When presented in a log2 scale, positive values correspond to enrichment, and negative values to depletion at UV sites. Each dot corresponds to one cell. Triangles represent the mean values of each independent experiment. a.u., arbitrary units. Statistical analysis is performed with a two-sided Student t-test with Welch’s correction (A, D, G-J) and by one-way ANOVA for multiple comparisons (B, G).

### Supplementary Figure S3

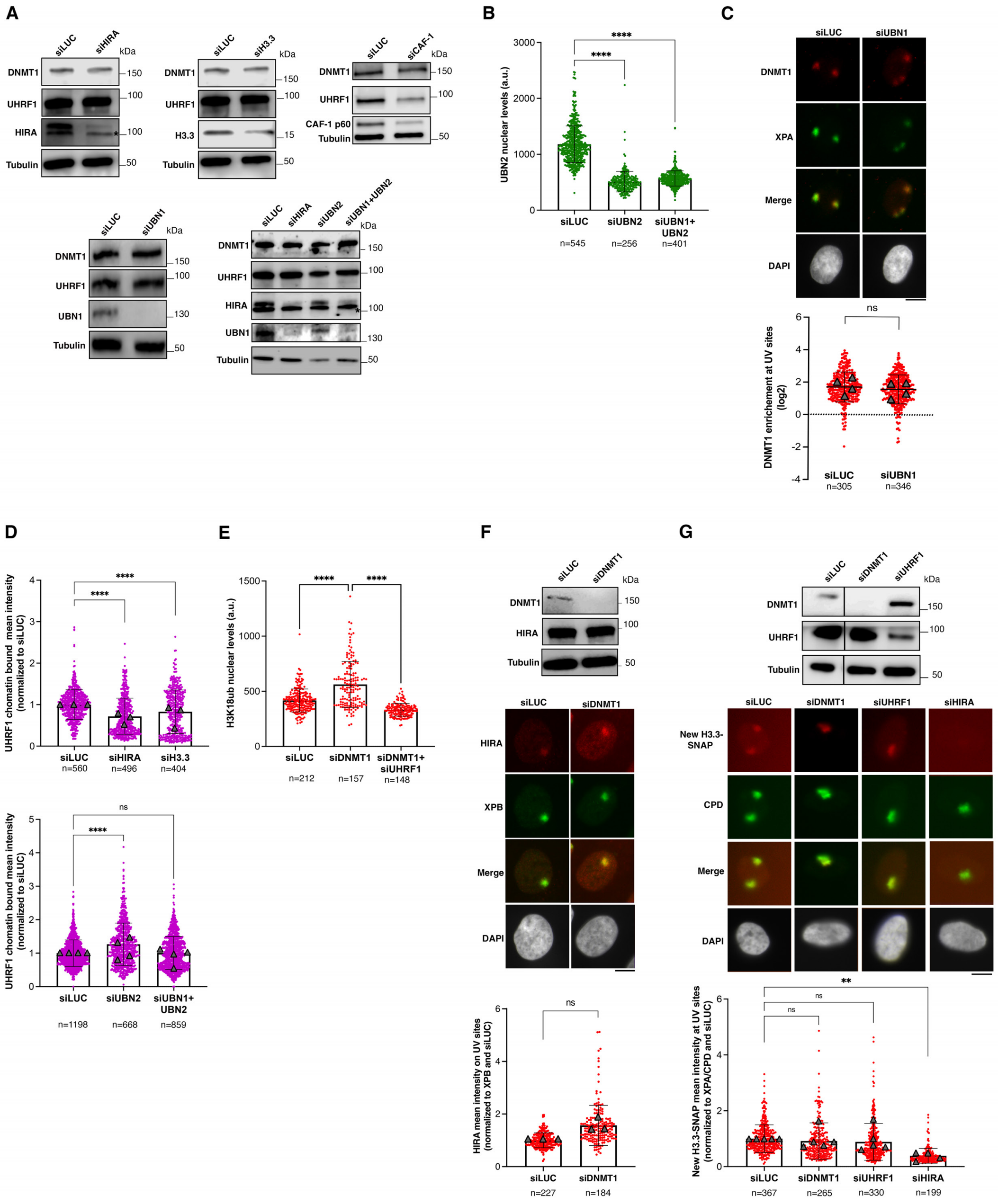

**Supplementary Figure S3: Crosstalk of the DNA methylation machinery with new H3.3 deposition during UV damage repair.**

(**A**) Western blot on total cell extracts from U2OS cells treated with the indicated siRNAs (siLUC, control; Tubulin, loading control). The asterisk indicates a non-specific band.

(**B**) UBN2 nuclear levels analyzed by immunofluorescence in U2OS cells treated with the indicated siRNAs (siLUC, control).

(**C**) Recruitment of DNMT1 (red) to UV damage sites (green) analyzed by immunofluorescence 2 h after local UVC irradiation in U2OS cells treated with the indicated siRNAs (siLuc, control).

(**D**) Quantification of UHRF1 chromatin-bound levels by immunofluorescence in U2OS cells treated with the indicated siRNAs (siLUC, control).

(**E**) Validation of H3K18ub antibody by immunofluorescence in U2OS cells treated with the indicated siRNAs (siLUC, control).

(**F**) Quantification of HIRA mean intensities at UV sites (marked by XPA or XPB) 2 h after local UVC irradiation in U2OS cells treated with siDNMT1 (siLUC, control). Results are normalized on the damage signal (XPA or XPB levels at UV sites). The efficiency of DNMT1 depletion is controlled by western blot (Tubulin, loading control).

(**G**) Quantification of new H3.3 deposition at UV damage sites (CPD) by immunofluorescence 1 h after local UVC irradiation in U2OS H3-SNAP cells treated with the indicated siRNAs (siLUC, control). Results are normalized on the damage signal (CPD or XPA levels at UV sites). Knock-down efficiencies are controlled by western blot (Tubulin, loading control).

Scale bars, 10 µm. Data on the graphs are presented as mean values ± SD from n cells scored in 2 to 4 independent experiments. Triangles represent the mean values of each independent experiment. Statistical analysis is performed by one-way ANOVA (B, E, G), Kruskal-Wallis test (D) or by two-sided Student’s t-test with Welch’s correction (C, F). a.u.: arbitrary units.

### Supplementary Figure S4

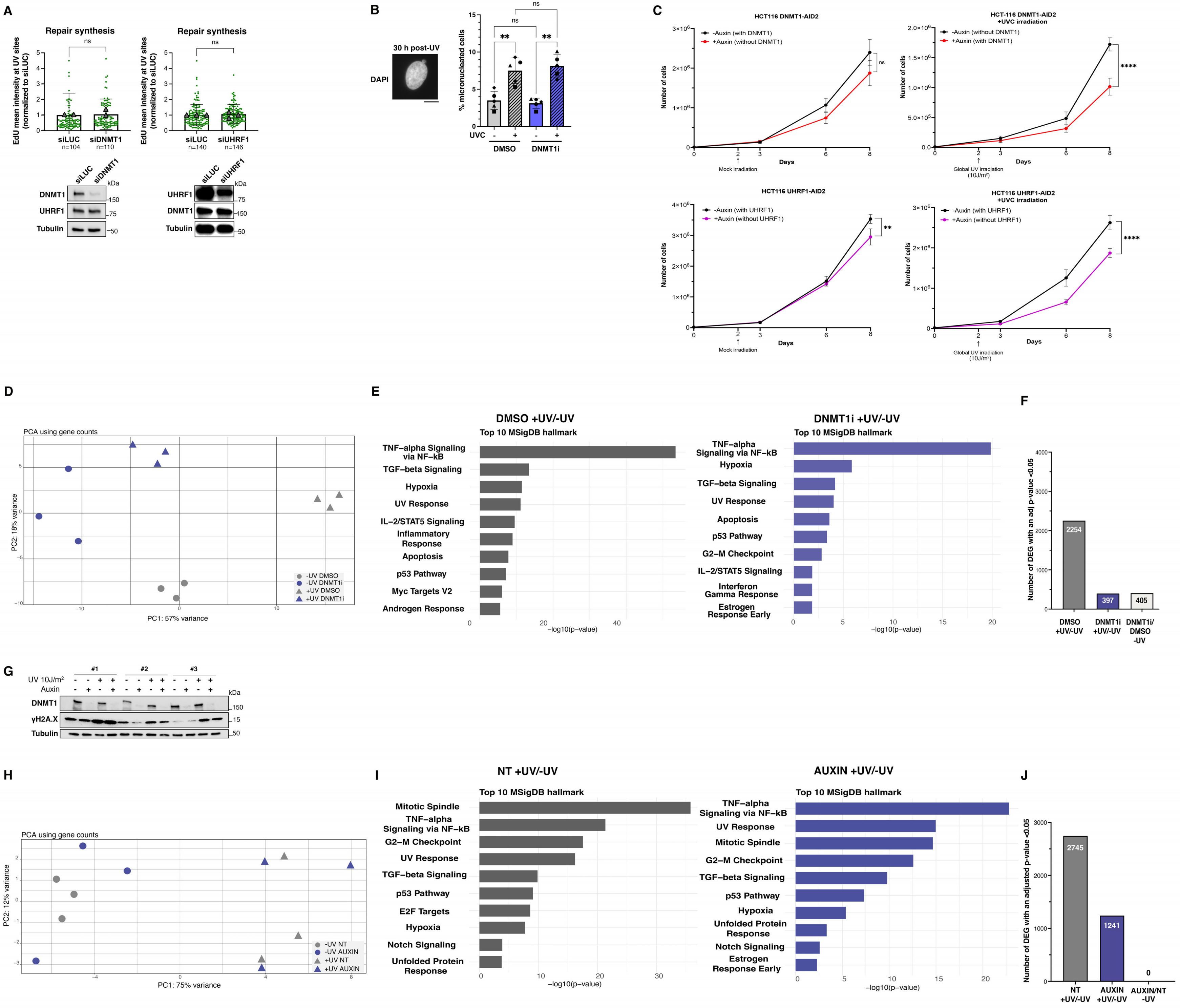

**Supplementary Figure S4: Functional relevance of DNA methylation maintenance during UV damage repair**

**(A)** Quantification of EdU levels at sites of local UVC irradiation in U2OS cells treated with the indicated siRNAs (siLUC, control). The efficiency of DNMT1 and UHRF1 knockdowns is controlled by western blot (Tubulin, loading control). The graphs show mean values ± SD from n cells scored in 2 or 3 independent experiments. Triangles represent the mean values of each independent experiment.

(**B**) Percentage of micronucleated U2OS cells (as shown on the example image, scale bar 10 μm) before and 30 h after global UVC irradiation in control conditions (DMSO) or upon treatment with a DNMT1 inhibitor around the time of irradiation (DNMT1i). Mean values ± SD from 5 independent experiments scoring at least 800 cells per condition.

(**C**) Proliferation assays in HCT116-DNMT1-AID2 and HCT116-UHRF1-AID2 cells performed in non-irradiated control cells (left) and in UV-irradiated cells (right). Mean values ± SEM from 4 (DNMT1) and 5 (UHRF1) independent experiments. Statistical analysis is performed by non-linear regression with a second order polynomial model.

(**D**) Principal component analysis of three replicate RNA-seq experiments in U2OS cells.

(**E**) Top 10 enriched pathways according to the Hallmark 2020 gene set of the Molecular Signature database. Gene ontology analysis was performed on Differentially Expressed Genes 2 h after global UV irradiation (adj. p-value <0.05) in U2OS cells treated with DMSO (control, left) or with DNMT1 inhibitor (right).

(**F**) Histogram showing the number of Differentially Expressed Genes (adj. p-value <0.05) in different pairwise comparisons 2 h after global UV irradiation in U2OS cells.

(**G**) Western blot on HCT116-DNMT1-AID2 total cell extracts to control DNMT1 degradation post Auxin addition and DNA damage levels (γH2A.X) after UV irradiation in three replicate RNA-seq experiments (Tubulin, loading control).

(**H**) Principal component analysis of three replicate RNA-seq experiments in HCT116-DNMT1-AID2 cells.

(**I**) Top 10 enriched pathways according to the Molecular Signature database and the KEGG pathway database. Gene ontology analysis was performed on Differentially Expressed Genes 2 h after global UV irradiation (adj. p-value <0.05) in HCT116-DNMT1-AID2 control cells (NT, left) or treated with auxin (AUXIN, right).

(**J**) Histogram showing the number of Differentially Expressed Genes (adj. p-value <0.05) in different pairwise comparisons 2 h after global UV irradiation in HCT116-DNMT1-AID2 cells.

**Supplementary Table S1: siRNA sequences**

| siRNA name | Target sequence (5' → 3') |
|---|---|
| siCAF-1 (p60) | AATCTTGCTCGTCATACCAAA |
| siDNMT1 | CGATGAGGAAGTCGATGATAA |
| siDNMT3A | GCATCCACTGTGAATGATAAG |
| siDNMT3B | AGGTTAATGCTGAAAAATC |
| siERCC1 | GACGUAAUUCCCGACUAU |
| siH3.3 | 1:1 combination of CUACAAAAGCCGCUCGCAA (H3.3A) and GCUAAGAGAGUCACCAUCAU (H3.3B) |
| siHIRA | GCUAAGAGAGUCACCAUCAU |
| siLIG1 | GGCATGATCCTGAAGCAGA |
| siLUC | CGTACGCGGAATACTTCGA |
| siUBN1 | AUACAGGACUUGAUCGAUA |
| siUBN2 | UCCUGUUCCUCCUCACUUA |
| siUHRF1 | GCCTTTGATTCGTTCCTTC |
| siXPF | GUAGGAUACUUGUGGUUGA |
| siXPG | GAAAGAAGAUGCUAAACGU |

**Supplementary Table S2: Plasmids**

| Plasmids | Construct details | Reference |
|---|---|---|
| MBD-mCherry | mCherry tag was inserted instead of GFP in MBD-GFP plasmid using AgeI and SacI restriction sites | [62] |
| UHRF1-WT-pEGFP | Human UHRF1 was inserted into peGFP-C2 plasmid using HindIII/KpnI | [11] |
| UHRF1-PHD-pEGFP | Plasmid encoding the PHD domain (aa 305-371) of human UHRF1 with N-terminal GFP-tag under the control of a CMV promoter was cloned using the Gibson technology | [11] |
| UHRF1-SRA-pEGFP | Plasmid encoding the SRA domain (aa 416-591) of human UHRF1 with N-terminal GFP-tag under the control of a CMV promoter was cloned using the Gibson technology | [11] |
| UHRF1-TTD-pEGFP | Plasmid encoding the TTD domain (aa 116-283) of human UHRF1 with N-terminal GFP-tag under the control of a CMV promoter was cloned using the Gibson technology | [11] |
| UHRF1-UBL-pEGFP | Plasmid encoding the UBL domain (aa 1-83) of human UHRF1 with N-terminal GFP-tag under the control of a CMV promoter was cloned using the Gibson technology | [11] |

**Supplementary Table S3: HCl denaturation conditions for 5mC and 5hmC immunodetection**

| Cell line | Antibody | HCl concentration | Time of exposure | Temperature |
|---|---|---|---|---|
| U2OS | 5mC | 2N | 20 minutes | Room Temperature |
| | 5hmC | 4N | 10 minutes | Room Temperature |
| NIH/3T3 | 5mC | 4N | 20 minutes | 37°C |
| | 5hmC | 4N | 30 minutes | Room Temperature |

**Supplementary Table S4: Antibodies**

| **Type** | **Antibody Target** | **Dilution** | **Species** | **Supplier(Reference)** | **Application** |
|---|---|---|---|---|---|
| Primary antibodies | 5hmC | 1:500 | Rabbit | Active motif (39770) | IF |
| | 5mC | 1:100 | Mouse | Abcam (Ab10805) | IF |
| | ATF-3 | 1:250 | Mouse | Abcam (Ab207434) | WB |
| | CPD | 1:1000 | Mouse | Kamiya Biomedical (MC-062) | IF |
| | CAF-1 p60 | 1:1000 | Mouse | Active Motif (39995) | WB |
| | DNMT1 (human) | 1:100/1:1000 | Rabbit | Abcam (ab188453) | IF/WB |
| | DNMT1 (mouse) | 1:1000 | Rabbit | Abcam (ab876549) | IF |
| | DNMT3A | 1:100 | Rabbit | Cell Signaling Technology (3598S) | IF |
| | DNMT3B | 1:500 | Rabbit | Cell Signaling Technology (67259S) | IF |
| | ERCC1 | 1:200/1:500 | Mouse | Santa Cruz Biotechnology (sc-17809) | IF/WB |
| | GFP | 1:50 | Rat | Santa Cruz Biotechnology (sc-101536) | IF |
| | H3K9me1 | 1:200 | Rabbit | Abcam (ab9045) | IF |
| | H3K9me2 | 1:250 | Rabbit | Diagenode (C15410060) | IF |
| | H3K9me3 | 1:500 | Rabbit | Active Motif (39765) | IF |
| | H3K18ub | 1:100 | Rabbit | Cell Signaling Technology (37771T) | IF |

| | | | | | |
|---|---|---|---|---|---|
| | H3K36me2 | 1:100/1:1000 | Rabbit | Cell Signaling Technology (2901T) | IF/WB |
| | HIRA | 1:100 | Mouse | Active Motif (39557) | IF/WB |
| | Tubulin | 1:10000 | Mouse | Sigma Aldrich (T9026) | WB |
| | UBN1 | 1:2000 | Rabbit | Abcam (ab101282) | WB |
| | UBN2 | 1:100 | Rabbit | Antibodies-Online (ABIN5999135) | IF |
| | UHRF1 (human and mouse) | 1:400/1:1000 | Mouse | Sigma Aldrich (MABE308) | IF/WB |
| | XPA | 1:400 | Mouse | BD Biosciences (556453) | IF |
| | XPB | 1:500 | Rabbit | Sigma Aldrich (XO879) | IF |
| | XPF | 1:100/1:1000 | Rabbit | Bethyl Laboratories (A301-315A) | IF/WB |
| | XPG | 1:500 | Rabbit | Bethyl Laboratories (A301-484A) | WB |
| | γH2A.X | 1:1000 | Rabbit | Cell Signaling Technology (2577s) | WB |
| Secondary Antibodies | Mouse HRP | 1:10000 | Rat | Tebu-bio (18-8817-33) | WB |
| | Rabbit HRP | 1:10000 | Mouse | Tebu-bio (18-8816-33) | WB |
| | Mouse Alexa Fluor 488 | 1:1000 | Goat | Invitrogen (A11029) | IF |
| | Mouse Alexa Fluor 568 | 1:1000 | Goat | Invitrogen (A11037) | IF |
| | Mouse Alexa Fluor 594 | 1:1000 | Goat | Invitrogen (A11032) | IF |
| | Mouse Alexa Fluor 647 | 1:1000 | Goat | Invitrogen (A21236) | IF |
| | Rabbit Alexa Fluor 488 | 1:1000 | Goat | Invitrogen (A11034) | IF |
| | Rabbit Alexa Fluor 568 | 1:1000 | Goat | Invitrogen (A11029) | IF |
| | Rabbit Alexa Fluor 594 | 1:1000 | Goat | Invitrogen (A11037) | IF |
| | Rabbit Alexa Fluor 647 | 1:1000 | Goat | Invitrogen (A21245) | IF |

**Supplementary Table S5: Genomic regions selected for adaptive sampling**

| Chromosome | Start coordinate | End coordinate | Region of interest |
|---|---|---|---|
| Chr2 | 176560000 | 176690000 | HOXD |
| Chr7 | 27210000 | 27360000 | HOXA |
| Chr12 | 53890000 | 54040000 | HOXC |
| Chr14 | 2100000 | 2500000 | RNA45S2 |
| Chr17 | 0 | 23345000 | chr17L |
| Chr17 | 27800000 | 84276897 | chr17R |
